# Insulin signaling regulates *Pink1* mRNA localization via modulation of AMPK activity to support PINK1 function in neurons

**DOI:** 10.1101/2023.02.06.527276

**Authors:** J. Tabitha Hees, Angelika B. Harbauer

**Affiliations:** TUM Medical Graduate Center, Technical University of Munich, Munich, Germany; Max Planck Institute for Biological Intelligence, Martinsried, Germany; Technical University of Munich, Institute of Neuronal Cell Biology, Munich, Germany; Munich Cluster for Systems Neurology, Munich, Germany

## Abstract

Mitochondrial quality control failure is frequently observed in neurodegenerative diseases. The detection of damaged mitochondria by stabilization of PTEN-induced kinase 1 (PINK1) requires transport of *Pink1* mRNA by tethering it to the mitochondrial surface. Here, we report that inhibition of AMPK by activation of the insulin signaling cascade prevents *Pink1* mRNA binding to mitochondria. Mechanistically, AMPK phosphorylates the RNA anchor complex subunit SYNJ2BP within its PDZ domain, a phosphorylation site that is necessary for its interaction with the RNA-binding protein SYNJ2. Interestingly, loss of mitochondrial *Pink1* mRNA association upon insulin addition is required for PINK1 protein activation and its function as a ubiquitin kinase in the mitophagy pathway, thus placing PINK1 function under metabolic control. Induction of insulin-resistance *in vitro* by the key genetic Alzheimer-risk factor apolipoprotein E4 retains *Pink1* mRNA at the mitochondria and prevents proper PINK1 activity especially in neurites. Our results thus identify a metabolic switch controlling *Pink1* mRNA localization and PINK1 activity via insulin and AMPK signaling in neurons and propose a mechanistic connection between insulin resistance and mitochondrial dysfunction.

## Introduction

Neurons are particularly challenged in maintaining and distributing mitochondria throughout neurites due to their highly extended and complex structures^1, 2^. Transport of nuclear-encoded mRNAs for mitochondrial proteins and their local translation support the upkeep of mitochondrial functionality within neuronal processes^3–5^. We have recently identified a neuron-specific mechanism that tethers the transcript encoding for the mitochondrial protein PTEN-induced kinase 1 (PINK1) and potentially also other nuclear-encoded mitochondrial transcripts to mitochondria thereby facilitating their transport and local translation in axons^5^. PINK1, mutated in certain familial forms of Parkinson’s disease (PD)^6^, is a short-lived protein^7^ that functions as a sensor for mitochondrial damage. Under physiological conditions, PINK1 is imported into mitochondria, cleaved and degraded^8^. In damaged mitochondria, however, the depolarized membrane potential impairs mitochondrial PINK1 import. Instead, PINK1 stabilizes on the outer mitochondrial membrane^9, 10^ and phosphorylates ubiquitin molecules at serine 65 leading to recruitment and partial activation of the E3 ubiquitin ligase Parkin^11–13^. PINK1 phosphorylates Parkin at serine 65 resulting in full activation of Parkin^14, 15^, which then ubiquitinates several proteins on the outer mitochondrial membrane. The ubiquitin molecules, in turn, are further phosphorylated by PINK1. The phosphorylated ubiquitin chains covering the damaged mitochondria are then recognized by autophagy receptors such as optineurin. This ultimately results in the formation of autophagosomes and their delivery to degradative lysosomes^16^. The interaction between *Pink1* mRNA and mitochondria is maintained by an anchoring complex comprising the outer mitochondrial membrane protein Synaptojanin 2 binding protein (SYNJ2BP, also called OMP25) and the phosphatidylinositol phosphatase Synaptojanin 2 (SYNJ2), which contains an RNA-binding motif^17^. SYNJ2BP is ubiquitously expressed and localized to the outer mitochondrial membrane due to its C-terminal transmembrane domain^18^. SYNJ2BP has a PDZ domain at its N-terminus, which specifically binds to a unique motif in the C-terminus of SYNJ2a^18^, a splice form of SYNJ2 that is primarily expressed in neurons^5^. SYNJ2a binds the *Pink1* mRNA, whereas SYNJ2BP serves as mitochondrial anchor tethering SYNJ2a and *Pink1* mRNA to mitochondria, ensuring a constant supply of fresh PINK1 protein by local translation in axons and dendrites^5^. This is required to support local mitophagy in axons in order to eliminate acutely damaged mitochondria^5, 19^. However, it still remains to be elucidated how localization of *Pink1* mRNA as well as PINK1 activation and function are regulated in response to local signaling pathways in neurons.

AMP-activated protein kinase (AMPK) is known as the master regulator of energy sensing and consequently closely associated with mitochondrial homeostasis^20^. AMPK is composed of three subunits: one catalytic α subunit and two regulatory β and γ subunits^21^. It has more than a hundred known targets including multiple proteins that are involved in various aspects of mitochondrial homeostasis such as mitochondrial biogenesis, mitochondrial fission and autophagy^20^, however, it has not been tied directly to PINK1/Parkin-dependent mitophagy. AMPK is generally activated in response to an increased AMP/ATP ratio, but also regulated by several upstream kinases, including AKT downstream of insulin receptor (IR) signalling. In response to the hormone insulin, the IR autophosphorylates and recruits IR substrate (IRS) adaptor proteins^22, 23^. Subsequently, phosphatidylinositol 3-kinase (PI3K) gets activated, thereby inducing AKT activity. AKT in turn inhibits AMPK by phosphorylating the catalytic α subunit^24–30^. Insulin signaling has also been shown to regulate neuronal mitochondrial function including ATP production, respiration, calcium buffering as well as protein biogenesis and homeostasis^31^. Interestingly, insulin resistance in the brain is associated with mitochondrial dysfunction^31^ and fittingly, type 2 diabetes represents a risk factor for both PD and Alzheimer’s disease (AD)^32, 33^. The strongest genetic risk factor for AD is the presence of the ε4 allele of the apolipoprotein E (ApoE)^34, 35^, which has mechanistically been linked to decreased insulin signaling and insulin resistance in the brain by trapping the IR in early endosomes^36^, yet how ApoE4 presence affects mitophagy remains to be determined.

In this study, we aimed to investigate how insulin and AMPK signaling regulate mitochondrial *Pink1* mRNA localization and PINK1 function. We identified SYNJ2BP as a novel substrate of AMPK downstream of insulin signaling. While SYNJ2BP phosphorylation is required for *Pink1* mRNA localization to mitochondria, insulin-induced AMPK inhibition and subsequent dissociation of *Pink1* mRNA from mitochondria promotes PINK1 protein activation. Disruption of insulin signaling by ApoE4 leads to impaired PINK1 activation and therefore mechanistically connects the frequently observed neurodegenerative phenotypes of insulin resistance and mitochondrial dysfunction.

## Results

### AMPK signaling regulates *Pink1* mRNA localization to mitochondria

In order to test whether mitochondrial localization of the *Pink1* mRNA is influenced by cellular metabolism, we treated mouse hippocampal neurons grown *in vitro* with either the AMP analogue 5-aminoimidazole-4-carboxamide ribonucleoside (AICAR) or Compound C (CC), which have been shown to activate and inhibit the master regulator kinase AMPK, respectively^37, 38^. To evaluate the impact of AICAR and CC on *Pink1* mRNA localization, we visualized mitochondria by expressing mitochondrially targeted mRaspberry and *Pink1* mRNA using the MS2/PP7-split Venus imaging that we established previously^5^. Briefly, this technique employs the high affinity interaction between two phage-derived capsid proteins (MS2 and PP7 coat proteins) and their respective RNA stem loops (MS2 and PP7), which we inserted downstream of the rat Pink1 3’UTR in 12 alternating copies. Co-expression of the so-tagged *Pink1* mRNA together with the coat proteins fused each to one half of split Venus allows the specific and background-free labelling of RNA in living cells^39^ (Extended Data Fig. 1a). While AICAR treatment had no effect on mitochondrial *Pink1* mRNA association, CC led to a loss of co-localization of *Pink1* mRNA with mitochondria in both the soma and neurites (Fig. 1a), which we quantified using the Manders’ coefficient (Fig. 1b,c). Importantly, the loss of co-localization was not due to a reduction of *Pink1* mRNA levels upon CC treatment (Extended Data Fig. 1b). We also determined the co-localization in a square within the cell body before and after 90-degree rotation of one imaging channel. The association between *Pink1* mRNA and mitochondria under control conditions was significantly higher than its corresponding rotated quantification, whereas the association in presence of CC was similar to chance level represented by its corresponding rotated quantification (Extended Data Fig. 1c). This indicates a CC specific effect on mitochondrial *Pink1* mRNA localization. Since CC is known to also affect other kinases^40^, we knocked down both isoforms of the catalytic subunit α of AMPK using short hairpin RNAs (as in^41^). Loss of AMPK activity by shRNA also reduced the mitochondrial localization of *Pink1* mRNA as measured in the soma (Fig. 1d,e), fully recapitulating the effect of CC. Thus, we identified AMPK to be a positive regulator of mitochondrial *Pink1* mRNA localization.

**Fig. 1:**
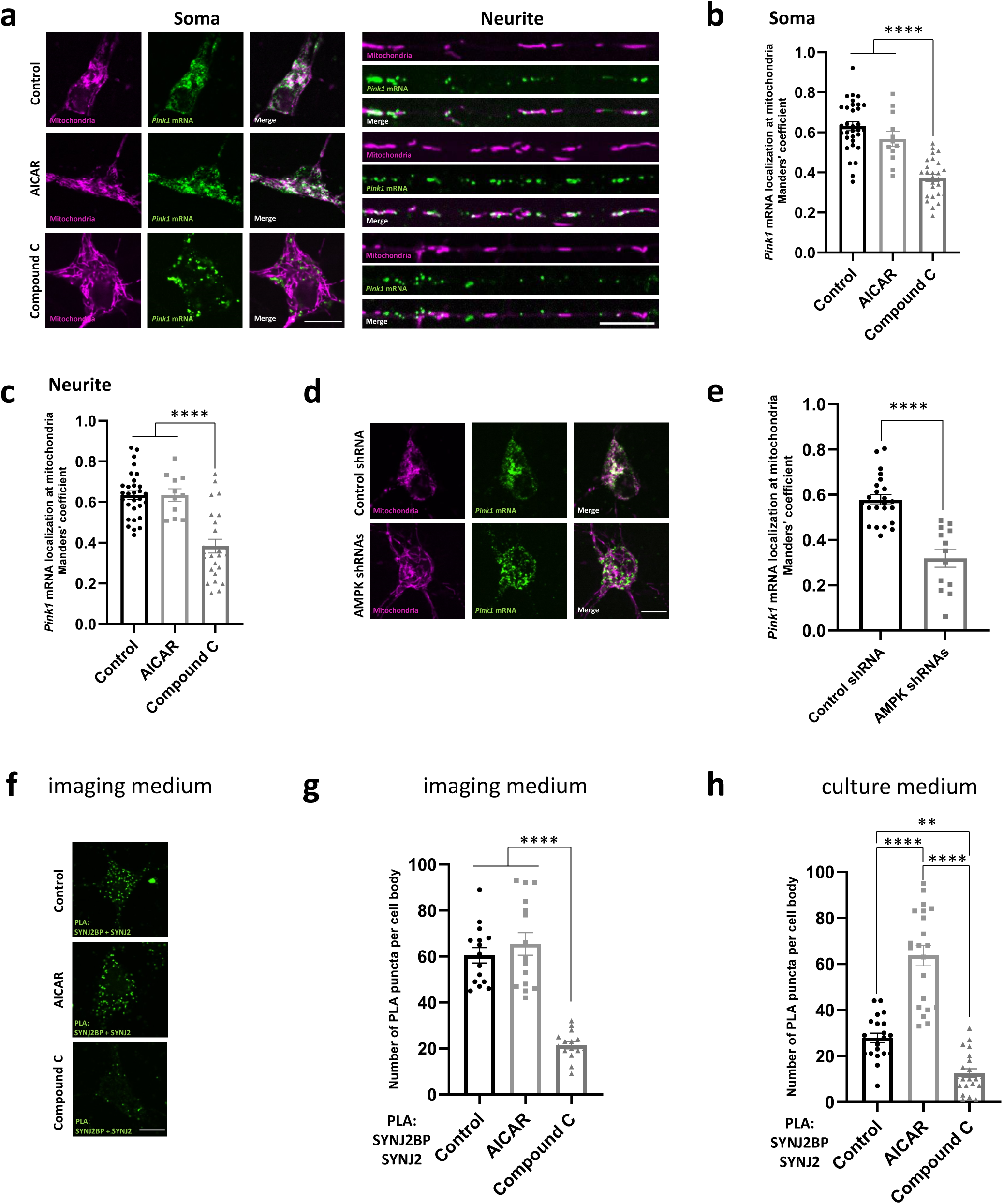
AMPK signaling regulates *Pink1* mRNA localization to mitochondria. **a** Representative images of *Pink1* mRNA visualized by the MS2/PP7 split Venus method and mitoRaspberry upon AMPK activation and inhibition using AICAR (1 mM, 2 h) and CC (20 µM, 2 h), respectively, in the soma and neurites. **b** Quantification of the Manders’ colocalization coefficient for the overlap between the *Pink1* mRNA and mitochondrial channel in the soma as in **a**. One-way ANOVA followed by Tukey’s post hoc test; n = 12-32; p < 0.0001 (****). **c** Quantification as in **b** for neurites. One-way ANOVA followed by Tukey’s post hoc test; n = 11-30; p < 0.0001 (****). **d** Representative images of *Pink1* mRNA and mitoRaspberry upon control or AMPK shRNA expression in the soma. **e** Quantification of the Manders’ colocalization coefficient for the overlap between the *Pink1* mRNA and mitochondrial channel as in **d**. Two-tailed student’s t-test; n = 13-23; p < 0.0001 (****). **f** Representative images of neuronal somas displaying the Proximity Ligation Assay (PLA) signal between SYNJ2BP and SYNJ2 upon control, AICAR (1 mM, 2 h) and CC (20 µM, 2 h) treatment in imaging medium (Hibernate E). **g** Number of PLA puncta per soma of neurons treated with the indicated drugs as in **f** in imaging medium (Hibernate E). One-way ANOVA followed by Tukey’s post hoc test; n = 15; p < 0.0001 (****). **h** Quantification as in **g** of neurons grown in full culture medium and treated with the indicated drugs. One-way ANOVA followed by Tukey’s post hoc test; n = 21; p < 0.01 (**), p < 0.0001 (****). All data are expressed as mean±SEM. All data points represent single cells coming from ≥3 biological replicates. Scale bars, 10 µm.

To better understand the mechanism, we investigated the proximity of the mitochondrial anchor SYNJ2BP to the RNA binding protein SYNJ2 by proximity ligation assay (PLA) using antibodies to detect the endogenous proteins^5^. In line with a loss of *Pink1* mRNA tethering to mitochondria, we observed a loss of proximity between SYNJ2 and SYNJ2BP upon CC treatment (Fig. 1f and Extended Data Fig. 1d), quantified by the somatic count of PLA dots (Fig. 1g). Interestingly, when we performed the experiment under similar conditions as our live cell imaging by using Hibernate E medium instead of B27-supplemented Neurobasal, we were not able to further increase the number of PLA dots by activation of AMPK using AICAR (Fig. 1f,g and Extended Data Fig. 1d), which is in line with the lack of effect of AICAR treatment on *Pink1* mRNA localization (Fig. 1a-c). However, when the experiment was performed in B27 containing culture medium instead of switching to Hibernate E imaging medium, the baseline number of PLA dots dropped and was now modulated in opposite directions by treatment with either AICAR or CC (Fig. 1h and Extended Data Fig. 1e). This suggests that the AMPK-dependent mechanism tethering *Pink1* mRNA to mitochondria responds to cues differentially present in both media.

### Insulin signaling regulates *Pink1* mRNA localization to mitochondria upstream of AMPK

One major signaling molecule absent in Hibernate E medium, but present in abundance in the neuronal supplement B27, is the peptide hormone insulin. Fittingly, activation of AKT downstream of insulin receptor (IR) signaling has already been shown to negatively regulate AMPK signaling^24–30^. We verified this negative regulation in neurons using lifetime imaging of a FRET-based AMPK activity sensor^42^. Indeed, if we added increasing amounts of insulin to the Hibernate E medium, we observed a dose-dependent decrease in AMPK activity, indicated by an increase in lifetime of the FRET donor (Extended Data Fig. 2a). This effect was prevented by co-treatment with inhibitors directed against the IR (GSK1904529A), PI3K (Wortmannin) or AKT (AKT inhibitor VIII) (Extended Data Fig. 2b), corroborating that insulin addition induces inhibition of AMPK activity via activation of PI3K and AKT also in neurons. Hence, we tested whether insulin treatment *per se* affects the localization of *Pink1* mRNA at mitochondria. Addition of insulin decreased the recruitment of *Pink1* mRNA to mitochondria both in the soma and in neurites as visualized by MS2/PP7-based mRNA imaging (Fig. 2a-c). This effect was dependent on the activity of the IR (Fig. 2a-c), as well as on the activity of PI3K and AKT (Extended Data Fig. 2c,d), corroborating the involvement of this signaling pathway in the regulation of mitochondrial *Pink1* mRNA association. This effect was also observed at lower insulin concentrations (Extended Data Fig. 2e) arguing that this scenario could happen under physiological conditions in the brain (low nanomolar concentrations)^43^. Also, the reduction in mitochondrial localization of the *Pink1* mRNA upon insulin addition was not as complete (down to chance levels; Extended Data Fig. 2f) as with CC treatment (Extended Data Fig. 1c), which is to be expected upon physiological inhibition of AMPK.

**Fig. 2:**
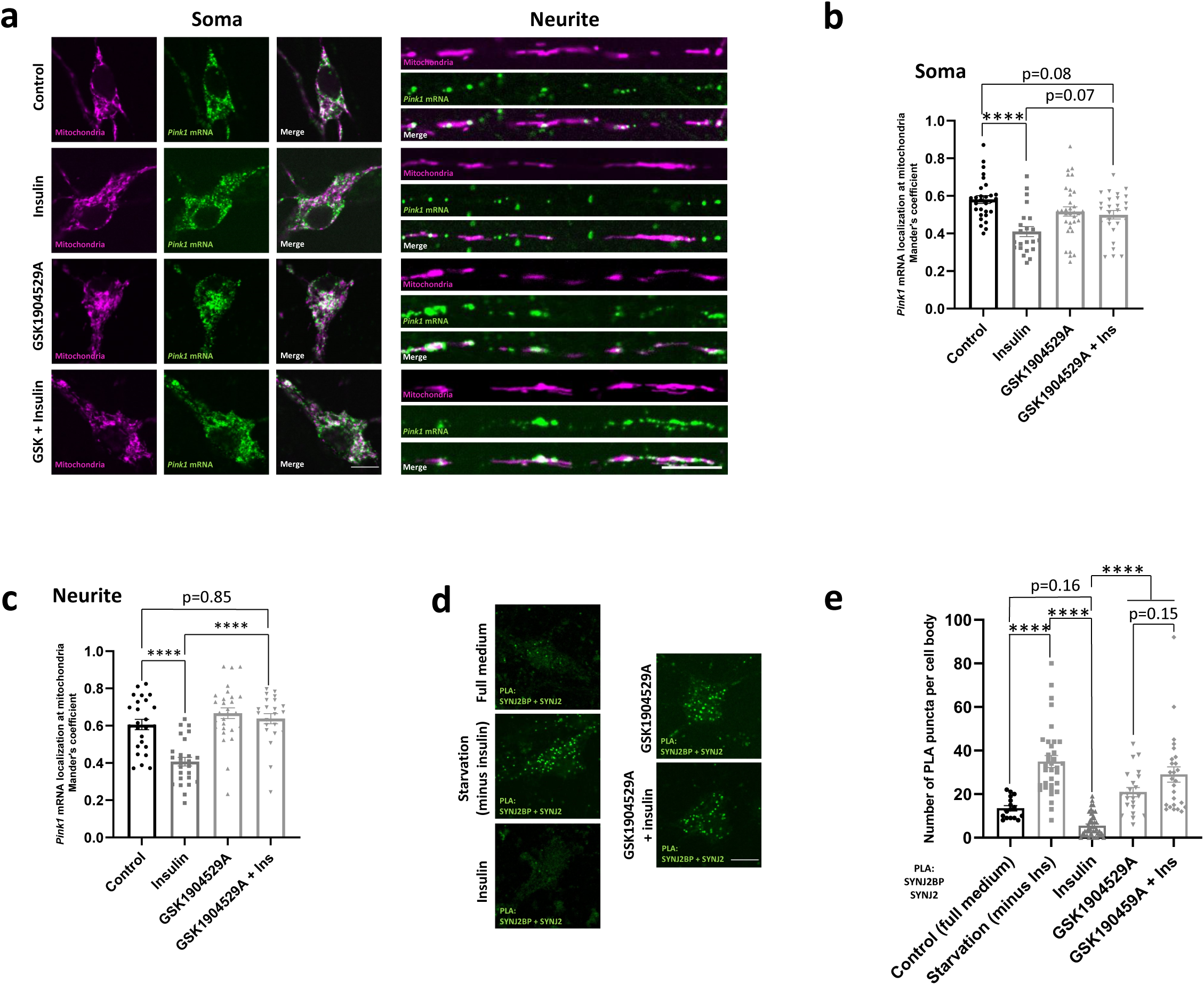
Insulin signaling regulates *Pink1* mRNA localization to mitochondria. **a** Representative images of *Pink1* mRNA visualized by the MS2/PP7 split Venus method and mitoRaspberry upon insulin (500 nM, 1 h) addition with or without pre-treatment with the IR inhibitor GSK1904529A (1 µM, 2 h) in the soma and neurites. **b** Quantification of the Manders’ colocalization coefficient for the overlap between the *Pink1* mRNA and mitochondrial channel in the soma. One-way ANOVA followed by Tukey’s post hoc test; n = 22-29; p < 0.0001 (****). **c** Quantification as in **b** for neurites. One-way ANOVA followed by Tukey’s post hoc test; n = 23-28; p < 0.0001 (****). **d** Representative images of neuronal somas analyzed by Proximity Ligation Assay (PLA) between SYNJ2BP and SYNJ2 in full medium upon insulin starvation and upon additional insulin (500 nM, 1 h) treatment with or without pre-treatment with the IR inhibitor GSK1904529A (1 µM, 2 h). **e** Number of PLA puncta per soma of neurons treated with the indicated drugs as in **d** in full medium or upon insulin withdrawal. One-way ANOVA followed by Tukey’s post hoc test; n = 16-45; p < 0.0001 (****). All data are expressed as mean±SEM. All data points represent single cells coming from ≥3 biological replicates. Scale bars, 10 µm.

Finally, we tested whether insulin also reduced the interaction between SYNJ2 and SYNJ2BP using PLA. As before, we observed a significant increase in proximity when neurons were cultured for two hours in medium lacking insulin (starvation). Addition of 500 nM insulin for one hour completely reversed this effect, but only when IR signaling was not inhibited (Fig. 2d,e and Extended Data Fig. 2g). Together, these results suggest that physiological inhibition of AMPK via IR signaling and AKT activation controls mitochondrial localization of *Pink1* mRNA by regulating the interaction of the RNA-binding protein SYNJ2 with its mitochondrial anchor SYNJ2BP.

### AMPK phosphorylates SYNJ2BP in its PDZ domain

Our observation that AMPK regulates the interaction between SYNJ2 and SYNJ2BP raised the question whether one of these proteins could be a direct substrate of AMPK. Interestingly, a phosphorylated peptide of SYNJ2BP matching the AMPK consensus motif^44^ (Fig. 3a) has been previously detected in high-throughput phospho-proteomics^45^. We therefore purified the cytosolic domain of rat SYNJ2BP from *E. Coli* and evaluated whether this protein could be phosphorylated *in vitro* by recombinant AMPK. To detect the phosphorylated protein, we employed the Phos-Tag approach^46^, which retards the electrophoretic mobility of phosphorylated proteins. We observed that the purified SYNJ2BP was already phosphorylated in *E. Coli*, which required pretreatment of the protein with recombinant calf intestinal phosphatase (CIP) before performing the kinase assay using AMPK (Extended Data Fig. 3a). We detected a slower migrating SYNJ2BP species upon addition of AMPK to the phosphorylation reaction, but this was prevented when the recombinant SYNJ2BP carried a mutation of the predicted phospho-site serine 21 to alanine (S21A, Fig. 3b). To test whether phosphorylation at this site responds to the physiological levels of AMPK activity present in neurons, we replaced the recombinant AMPK with lysates obtained from cortical neurons grown *in vitro* in the presence or absence of B27 (and therefore insulin), as well as lysates of neurons treated with AICAR to stimulate AMPK activity. Fittingly, we observed the slower migrating, phosphorylated SYNJ2BP only upon B27 starvation or AMPK activation, but not when neurons were grown in insulin containing medium or when the S21A mutant was used (Fig. 3c).

**Fig. 3:**
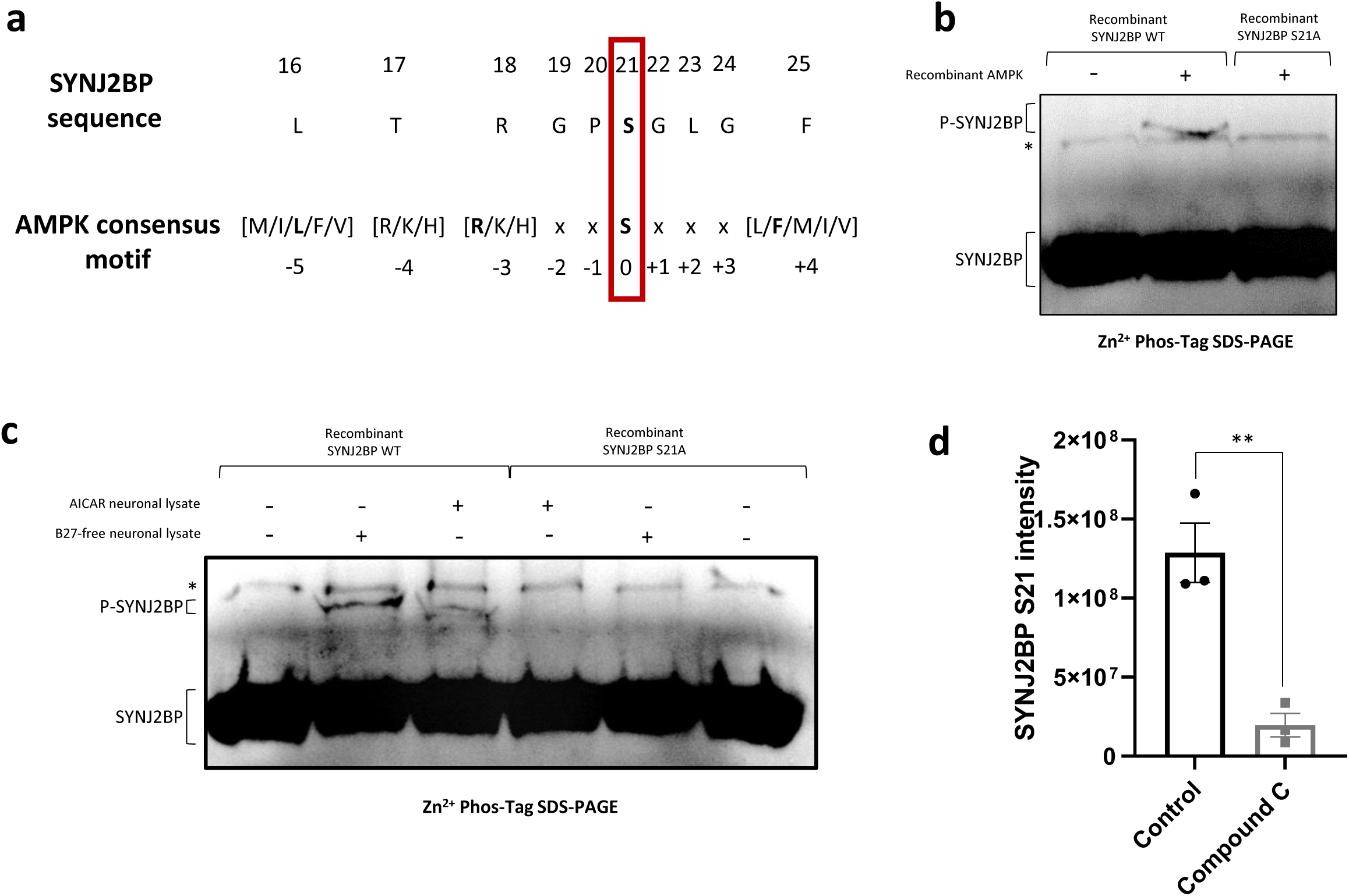
AMPK phosphorylates SYNJ2BP in its PDZ domain. **a** Schematic showing the SYNJ2BP sequence around its proposed phosphorylation site S21 and the AMPK consensus motif. **b** *In vitro* kinase assay analyzed on a Zn^2+^-Phos-Tag SDS-PAGE using recombinant AMPK as well as recombinant SYNJ2BP WT and S21A, respectively, and decorated with a SYNJ2BP antibody. Note the appearance of a slower migrating species only in the presence of AMPK and SYNJ2BP WT. The asterisk (*) denotes an unspecific band present in all samples. **c** *In vitro* phosphorylation assay as in **b**, using recombinant SYNJ2BP WT and S21A, respectively, treated with lysates from cortical neurons grown *in vitro* in the presence of AICAR or in the absence of the B27 supplement. Note the appearance of a slower migrating species only in the presence of AICAR-treated or B27-starved lysates and SYNJ2BP WT. The asterisk (*) denotes an unspecific band present in all samples. **d** Primary cortical neurons overexpressing myc-tagged SYNJ2BP WT by lentiviral transduction were cultured in insulin-free medium and treated with or without the AMPK inhibitor CC (20 µM, 2 h). The SYNJ2BP S21 intensity is shown upon phospho-peptide enrichment and LC MS/MS analysis. Two-tailed student’s t-test; n = 3; p < 0.01 (**). All data are expressed as mean±SEM. All data points represent biological replicates.

To test whether this phospho-site is observed in living neurons we employed phospho-peptide enrichment prior to mass spectrometry (phospho-MS). However, due to the low abundance of SYNJ2BP in cultured neurons (Extended Data Fig. 3b) we could not detect a phosphorylated peptide. 5 We therefore overexpressed myc-tagged SYNJ2BP in neurons by lentiviral transduction, which increased its cellular abundance (Extended Data Fig. 3c) and finally allowed us to observe a phosphorylated peptide upon phospho-MS, indeed showcasing phosphorylation at serine 21 (Extended Data Fig. 3d). Fittingly, SYNJ2BP serine 21 phosphorylation was reduced in cortical neurons that were grown in insulin-free medium for 2 h and at the same time treated with the AMPK inhibitor CC (Fig. 3d), while the total levels of SYNJ2BP did not decrease (Extended Data Fig. 3e). Together, these results indicate that SYNJ2BP is a substrate of AMPK.

### SYNJ2BP S21 phosphorylation regulates *Pink1* mRNA localization to mitochondria

Our observation of AMPK-mediated phosphorylation at serine 21 in SYNJ2BP prompted us to investigate whether mutating this residue to alanine or to the phospho-mimetic amino acid glutamate (S21E) would replicate the effects of AMPK on *Pink1* mRNA localization visualized by the MS2-PP7 split Venus imaging. Indeed, while expression of an shRNA resistant wild type (WT) version of SYNJ2BP was fully able to rescue the loss of mitochondrial localization of *Pink1* mRNA upon SYNJ2BP knock down, only the phospho-mimetic S21E, but not the phospho-ablative S21A mutant was also able to rescue *Pink1* transcript localization (Fig. 4a,b). Importantly, this was not due to reduced expression or stability of these rescue constructs (Extended Data Fig. 4a,b). This fully supports the model that AMPK-mediated phosphorylation at S21 of SYNJ2BP positively regulates *Pink1* mRNA recruitment to mitochondria.

**Fig. 4:**
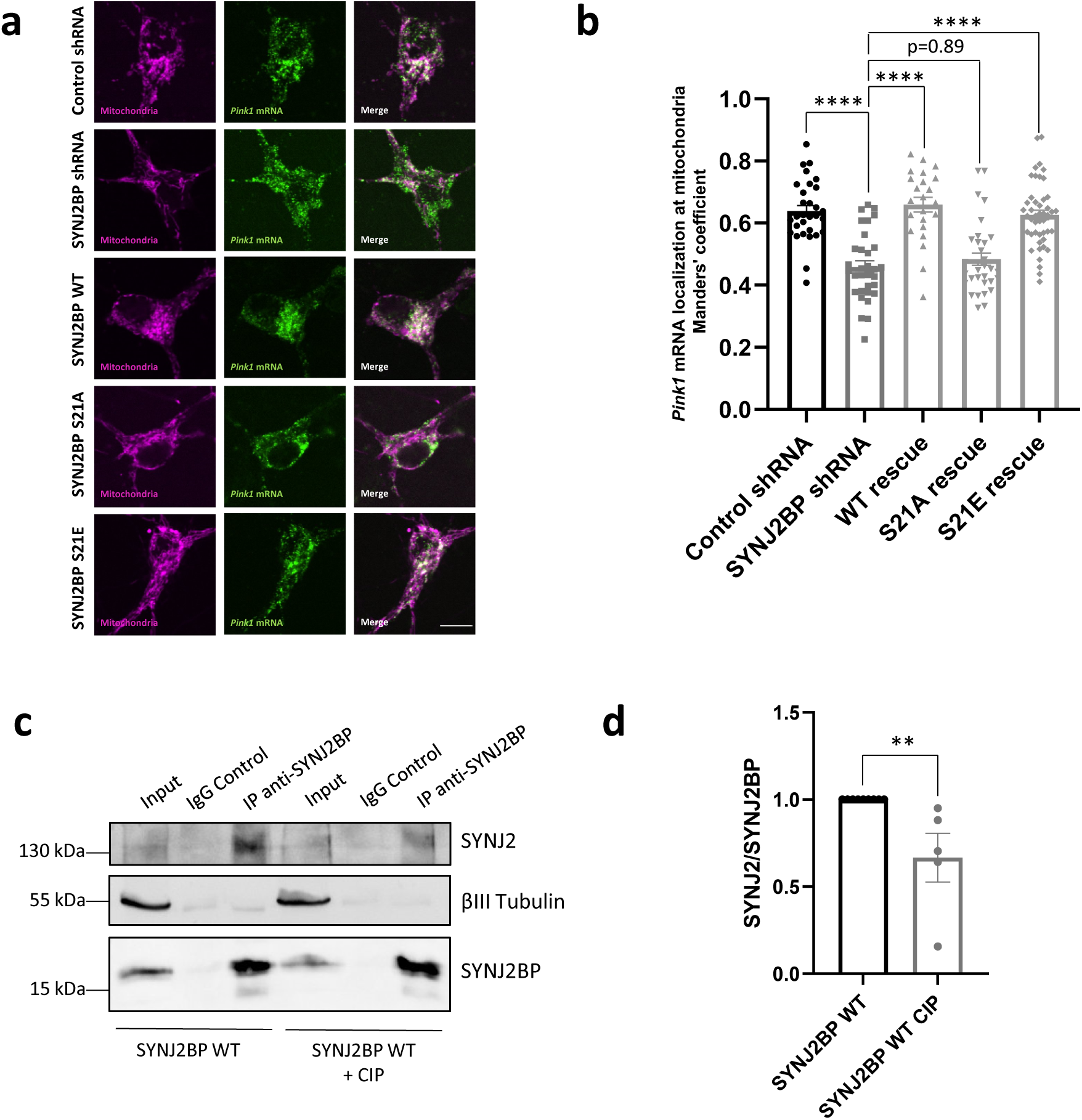
SYNJ2BP phosphorylation regulates *Pink1* mRNA localization to mitochondria. **a** Representative images of *Pink1* mRNA visualized by the MS2/PP7 split Venus method and mitoRaspberry upon control or SYNJ2BP shRNA expression combined with overexpression of SYNJ2BP WT, S21A or S21E in the soma. **b** Quantification of the Manders’ colocalization coefficient for the overlap between the *Pink1* mRNA and mitochondrial channel as in **a**. One-way ANOVA followed by Tukey’s post hoc test; n = 23-46; p < 0.0001 (****). **c** Representative immunoblot image of immunoprecipitation of SYNJ2BP using lysates of cortical neurons lentivirally overexpressing myc-tagged SYNJ2BP WT treated with and without calf intestinal phosphatase (CIP). Note, less SYNJ2 co-precipitated with SYNJ2BP from lysates treated with CIP. **d** Quantification of the SYNJ2 protein bands normalized to the respective SYNJ2BP bands as well as the input band as in **c**. Two-tailed student’s t-test, n = 5, p < 0.01 (**). All data are expressed as mean±SEM. Data points represent biological replicates (**d**) or single cells coming from 5 biological replicates (**b**). Scale bar, 10 µm.

To test whether the interaction of SYNJ2BP with SYNJ2 depends on its phosphorylation status, we immunoprecipitated virally overexpressed myc-tagged SYNJ2BP from cortical neurons grown in insulin-free medium for 2 h and determined the amount of co-isolated endogenous SYNJ2. We detected a specific interaction between the two proteins as an abundant neuronal protein, β-III-tubulin, was not co-isolated (Fig. 4c). This interaction was significantly diminished, when the lysates were treated with CIP (Fig. 4c,d). This shows that phosphorylation is a crucial prerequisite for efficient binding of SYNJ2 by SYNJ2BP and hence will regulate the localization of *Pink1* mRNA to mitochondria.

### Phospho-mimetic SYNJ2BP restores mitochondrial *Pink1* mRNA localization upon AMPK inhibition

As we have observed that the phospho-mimetic S21E mutant of SYNJ2BP was sufficient to restore *Pink1* mRNA localization (Fig. 4a,b), we wondered whether expression of this mutant could overcome the effects of AMPK inhibition. Therefore, we tested whether SYNJ2BP S21E overexpression would prevent the CC-mediated dissociation of *Pink1* mRNA from mitochondria, using the MS2-PP7 split Venus method. Indeed, the loss of co-localization induced by CC treatment was greatly diminished both in the soma and in neurites when instead of WT SYNJ2BP the phospho-mimetic S21E mutant was overexpressed (Fig. 5a,b and Extended Data Fig. 5a,b). Similarly, the effect of insulin addition was 6 abolished upon expression of SYNJ2BP S21E (Fig. 5c,d and Extended Data Fig. 5c,d), fully supporting the model that phosphorylation of SYNJ2BP S21 underlies the regulation of *Pink1* mRNA localization by insulin and AMPK signaling.

**Fig. 5:**
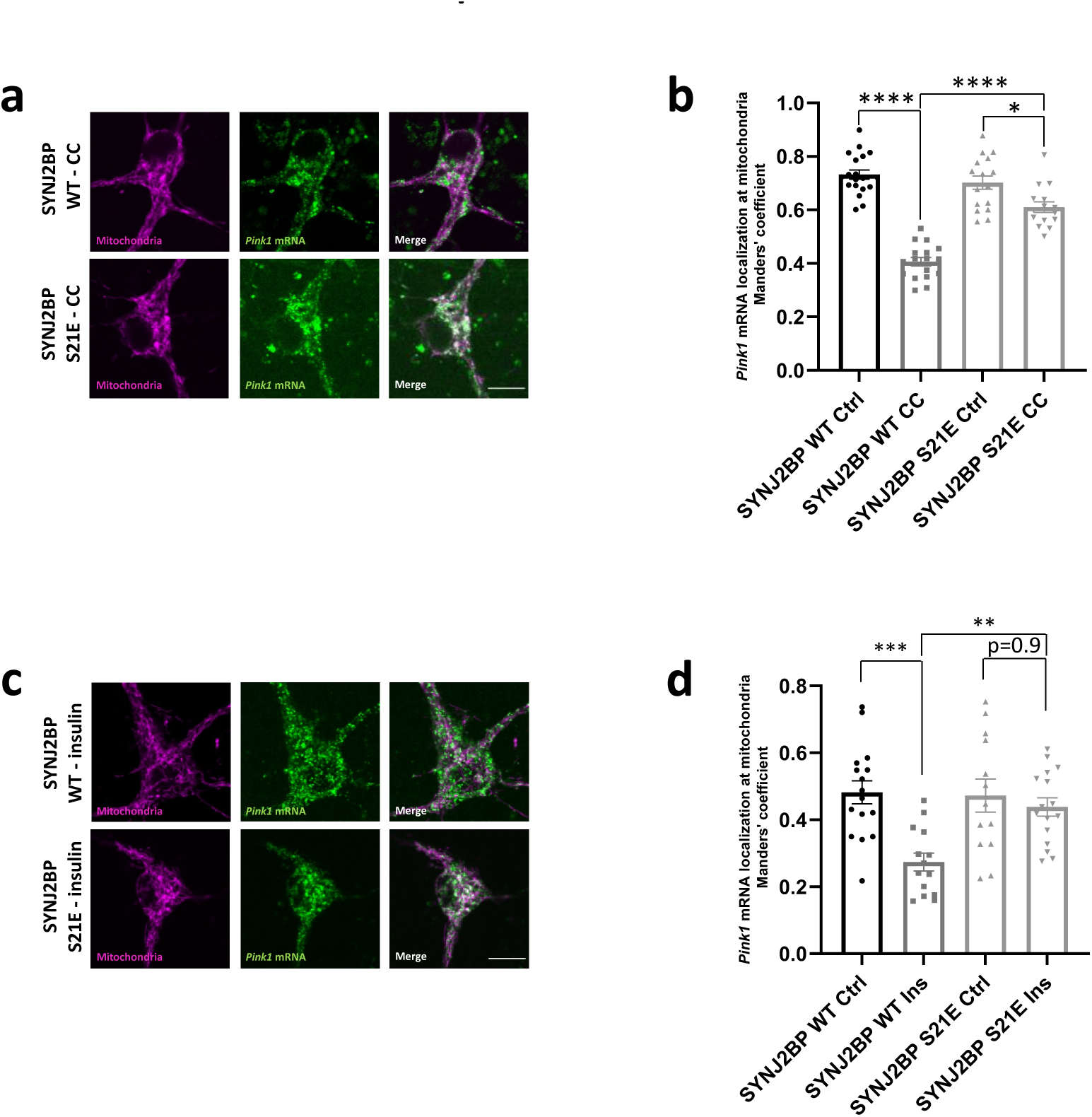
Phospho-mimetic SYNJ2BP restores mitochondrial *Pink1* mRNA localization upon AMPK inhibition. **a** Representative images of *Pink1* mRNA visualized by the MS2/PP7 split Venus method and mitoRaspberry upon CC (20 µM, 2 h) treatment combined with overexpression of SYNJ2BP WT and S21E, respectively. **b** Quantification of the Manders’ colocalization coefficient for the overlap between the *Pink1* mRNA and mitochondrial channel in the soma of neurons overexpressing SYNJ2BP WT or S21E and treated with or without CC (20 µM, 2 h). One-way ANOVA followed by Tukey’s post hoc test; n = 16-19; p < 0.05 (*), p < 0.0001 (****). **c** Representative images of *Pink1* mRNA and mitoRaspberry upon insulin (500 nM, 1 h) treatment combined with overexpression of SYNJ2BP WT and S21E, respectively. **d** Quantification of the Manders’ colocalization coefficient for the overlap between the *Pink1* mRNA and mitochondrial channel in the soma of neurons overexpressing SYNJ2BP WT or S21E and treated with or without insulin (500 nM, 1 h). One-way ANOVA followed by Tukey’s post hoc test; n = 13-16; p < 0.01 (**); p < 0.001 (***). All data are expressed as mean±SEM. All data points represent single cells coming from ≥3 biological replicates. Scale bars, 10 µm.

### Insulin supports PINK1 activation and mitophagy

The main function of PINK1 is the tagging of damaged mitochondria with phosphorylated ubiquitin, which is added by the recruitment of the E3-Ligase Parkin to the organelle. Therefore, we analyzed whether recruitment of Parkin to damaged mitochondria was affected by the presence of insulin using live cell microscopy. We took advantage of the decreased mitochondrial density in neurites to allow for the visualization of YFP-Parkin enrichment on mitochondria upon mitophagy induction. Unexpectedly, we observed that recruitment of Parkin to mitochondria upon mitochondrial damage with complex III inhibitor Antimycin A (AA) was decreased in neurites grown in the absence of insulin overnight (Fig. 6a,b), despite increased mRNA tethering (Fig. 2a-c). This suggested that mRNA tethering and mitophagy induction might be reciprocally regulated. We also tested the overall amount of mitophagy and acidification of the resulting autophagolysosomes as measured by the pH-sensitive excitation shift of mitochondrially targeted mKeima (mito-mKeima)^47^. We confirmed the positive effect of insulin signaling on mitophagy under mildly damaging conditions using this assay (5 nM AA, Fig. 6c,d). At higher concentrations of the mitochondrial toxin (20 µM AA), this difference disappeared (Extended Data Fig. 6a,b). As the mito-mKeima assay measures all mitophagy events and is not exclusive to PINK1-dependent mitophagy, this effect may be due to induction of compensatory mitophagy pathways under these conditions of extensive mitochondrial damage.

**Fig. 6:**
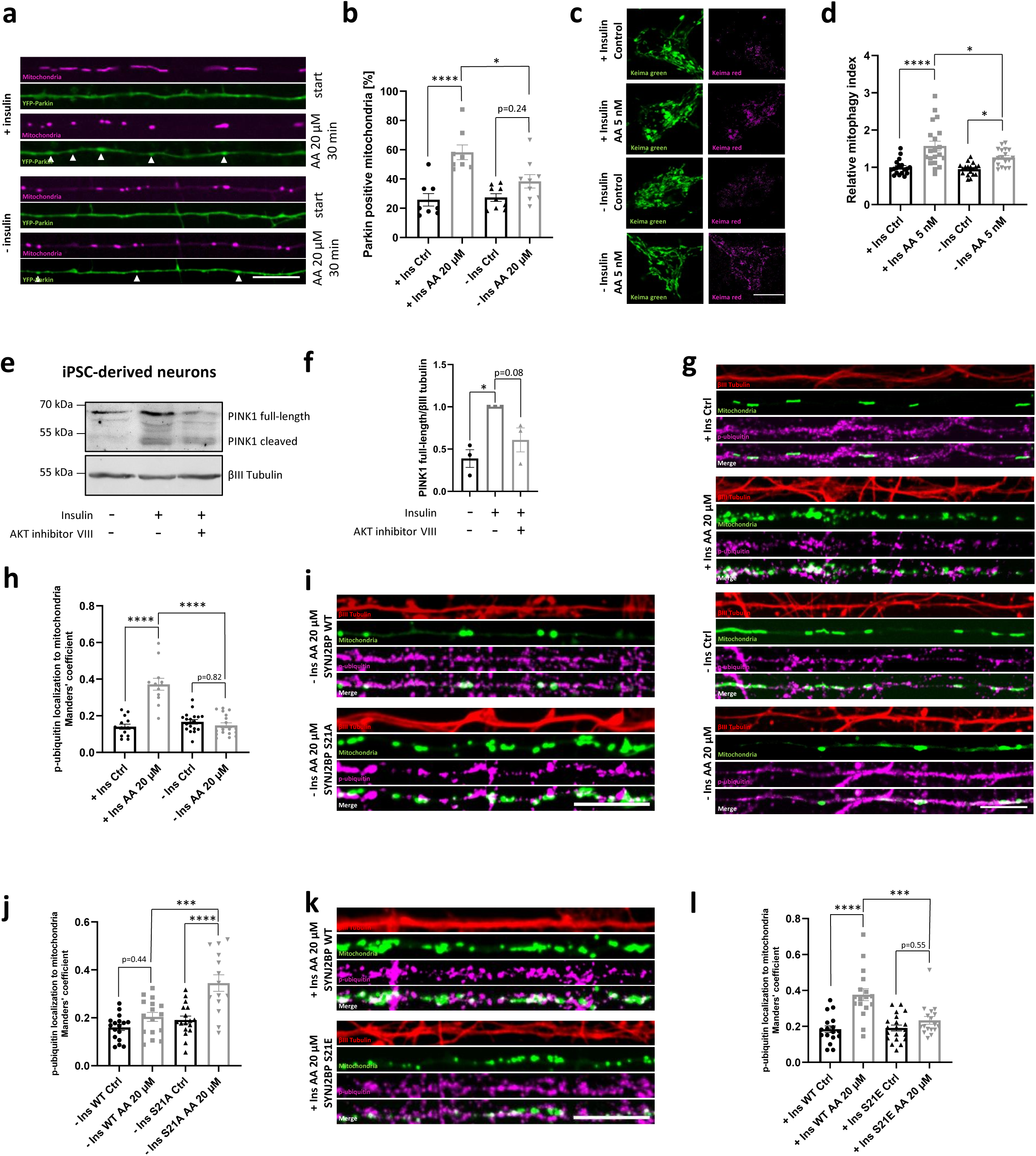
Insulin supports PINK1 activation and mitophagy. **a** Representative images of neurites overexpressing YFP-Parkin and mitoRaspberry cultured in the presence or absence of insulin overnight prior to treatment with 20 µM AA. The white arrowheads indicate Parkin recruitment to mitochondria monitored with live cell imaging before (start) and after 30 min of AA addition. **b** Quantification of mitochondria co-localizing with Parkin before and after AA treatment in presence or absence of insulin as in **a**. One-way ANOVA followed by Tukey’s post hoc test; n = 8-9; p < 0.05 (*), p < 0.0001 (****). **c** Representative images of neurons overexpressing the pH-sensitive fluorophore mito-mKeima cultured in the presence or absence of insulin overnight prior to treatment with or without 5 nM AA. **d** Quantification of the relative mitophagy index in neurons as in **c**. One-way ANOVA followed by Tukey’s post hoc test; n = 18-20; p < 0.05 (*), p < 0.0001 (****). **e** Representative immunoblot image of human iPSC-derived cortical neurons cultured in presence or absence of insulin (2 h) and treated with or without the AKT inhibitor VIII (10 µM, 2 h). **f** Quantification of the full-length PINK1 protein bands normalized to the βIII tubulin signal as in **e**. One-way ANOVA followed by Tukey’s post hoc test; n = 3; p < 0.05 (*). **g** Representative images of neurites overexpressing mito-meGFP cultured in the presence or absence of insulin overnight prior to treatment with 20 µM AA and stained with an antibody against phospho-ubiquitin (S65) and βIII tubulin. **h** Quantification of phospho-ubiquitin (S65) localization to mitochondria using the Manders’ colocalization coefficient as in **g**. One-way ANOVA followed by Tukey’s post hoc test; n = 12-19; p < 0.0001 (****). **i** Representative images of neurites overexpressing mito-meGFP as well as SYNJ2BP WT or S21A cultured in the absence of insulin overnight prior to treatment with 20 µM AA and stained with an antibody against phospho-ubiquitin (S65) and βIII tubulin. **j** Quantification of phospho-ubiquitin (S65) localization to mitochondria using the Manders’ colocalization coefficient. One-way ANOVA followed by Tukey’s post hoc test; n = 14-17; p < 0.001 (***), p < 0.0001 (****). **k** Representative images of neurites overexpressing mito-meGFP as well as SYNJ2BP WT or S21E cultured in the presence of insulin overnight prior to treatment with 20 µM AA and stained with an antibody against phospho-ubiquitin (S65) and βIII tubulin. **l** Quantification of phospho-ubiquitin (S65) localization to mitochondria using the Manders’ colocalization coefficient. One-way ANOVA followed by Tukey’s post hoc test; n = 17-20; p < 0.001 (***), p < 0.0001 (****). All data are expressed as mean±SEM. Data points represent biological replicates (**f**) or single cells coming from ≥2 biological replicates (**b**, **d**, **h**, **j** and **I**). Scale bars, 10 µm.

To understand the opposing effects of insulin on mRNA tethering and mitophagy induction, we sought to investigate its effect on PINK1 translation and activity. We took advantage of a PINK1 antibody recognizing the human PINK1 protein and compared the levels of PINK1 protein in human iPSC-derived cortical neurons cultured in medium either supplemented with or without insulin. Interestingly, already 2 h of insulin withdrawal reduced all major PINK1 protein species, as did AKT inhibition (Fig. 6e,f), in accordance with the very short half-life of the protein^7, 48, 49^ and with the reduced Parkin recruitment under these conditions (Fig. 6a,b). In HEK293 cells, inhibition of insulin signaling did not result in lower PINK1 expression (Extended Data Fig. 6c,d), suggesting that this is a neuron-specific mechanism and therefore might be tied to the neuron-specific tethering of the *Pink1* mRNA to mitochondria^5^. Hence, the mitophagy defect observed in the Parkin translocation assay upon insulin withdrawal (Fig. 6a,b) could be explained by a reduced availability of the *Pink1* transcript for translation due to its mitochondrial association (Fig. 2a-c).

To further test PINK1 activity, we probed the generation of phospho-ubiquitin at mitochondria using a phospho-specific antibody in primary mouse hippocampal neurons, cultured either in the presence or 7 absence of insulin overnight. While the baseline levels of mitochondrial phospho-ubiquitin remained unchanged, neurons grown in insulin-free media were unable to boost mitochondrial phospho-ubiquitination upon induction of mitophagy with AA (Fig. 6g,h), consistent with reduced PINK1 availability. This boost in PINK1 availability/activity did not depend on general translational regulation by mammalian target of rapamycin (mTOR) signaling, as mTOR inhibition using Torin-2 still resulted in mitochondrial phospho-ubiquitination upon AA treatment, but could be suppressed by addition of the AMPK activator AICAR (Extended Data Fig. 6e,f). This suggested that the activity of PINK1 is regulated by AMPK downstream of insulin signaling independently of mTOR signaling, and hence could be regulated at the level of SYNJ2BP phosphorylation. Therefore, we tested whether mutating SYNJ2BP S21 would be sufficient to prevent the block in PINK1 activity seen upon insulin withdrawal. Indeed, expression of the phospho-ablative SYNJ2BP S21A but not WT prevented the block in PINK1 activity in medium lacking insulin (Fig. 6i,j). This is consistent with the model that sequestering of the mRNA at mitochondria is responsible for the lack of PINK1 production, as the S21A mutant is unable to efficiently recruit the *Pink1* mRNA to mitochondria (Fig. 4a,b). Concurrently, in insulin containing medium, PINK1 activation can occur in the presence of WT SYNJ2BP, but is blocked in the presence of the phospho-mimetic S21E mutant (Fig. 6k,l), which retains the mRNA at mitochondria even in the presence of insulin (Fig. 5c,d). Thus, insulin-mediated uncoupling of the *Pink1* mRNA from mitochondria is a prerequisite for PINK1 activation and efficient induction of PINK1/Parkin-dependent mitophagy.

### ApoE4 inhibits insulin-regulated *Pink1* mRNA localization and PINK1 activation

Brain insulin resistance is believed to contribute to the observed metabolic changes in AD^50^, yet no direct connection between insulin resistance and mitochondrial dysfunction has been shown in neurons. Our results suggest that defective insulin signaling will have direct consequences on *Pink1* mRNA localization and downstream mitochondrial quality. Therefore, we tested whether *Pink1* mRNA localization is altered in an *in vitro* model of insulin resistance by application of human ApoE4, a highly prevalent AD risk factor^34, 35^. ApoE4 application to cultured neurons has previously been shown to blunt IR signaling by sequestration of the IR into endosomes^36^. We confirmed that application of ApoE4 also prevented the insulin-induced decrease in AMPK activity, using lifetime measurement of the FRET-based AMPK activity sensor. Indeed, neurons treated with ApoE4 overnight were unable to modulate AMPK activity in response to insulin application (Extended Data Fig. 7a). This effect was specific to ApoE4 because treatment with its homolog ApoE3 did not alter the response to insulin (Extended Data Fig. 7a). Furthermore, ApoE4 treatment prevented the insulin-mediated decline in mitochondrial *Pink1* mRNA localization both in the soma and in neurites visualized with the MS2/PP7-split Venus system. ApoE3 exposure, on the other hand, did not affect the insulin-induced impact on *Pink1* mRNA localization (Fig. 7a-c). In contrast to insulin, CC-induced loss of mitochondrial *Pink1* mRNA localization was not affected by ApoE4 treatment (Extended Data Fig. 7b,c), suggesting that the effect of ApoE4 on 8 *Pink1* mRNA localization is indeed upstream of AMPK-mediated phosphorylation of SYNJ2BP. Therefore, we tested whether overexpression of the phospho-ablative SYNJ2BP S21A mutant could prevent the ApoE4 effect in the presence of insulin. Indeed, mitochondrial *Pink1* mRNA localization was greatly diminished upon insulin and ApoE4 treatment when instead of WT SYNJ2BP the phospho-ablative S21A mutant was overexpressed (Fig. 7d,e). As we have seen that untethering of the *Pink1* mRNA from mitochondria is required for full PINK1 activity, we tested whether ApoE4 application inhibited the accumulation of phospho-ubiquitin on mitochondria upon mitochondrial intoxication with AA. In insulin containing medium, AA induced mitochondrial phospho-ubiquitin levels in neurites, but not when the cells were treated with ApoE4 overnight (Fig. 7f,g). Together, these data suggest that dysregulation of *Pink1* mRNA localization and PINK1 activity are contributing to mitochondrial dysfunction under pathological conditions modeling insulin resistance *in vitro*. This provides a mechanistic connection between these two epiphenomena commonly seen in AD.

**Fig. 7:**
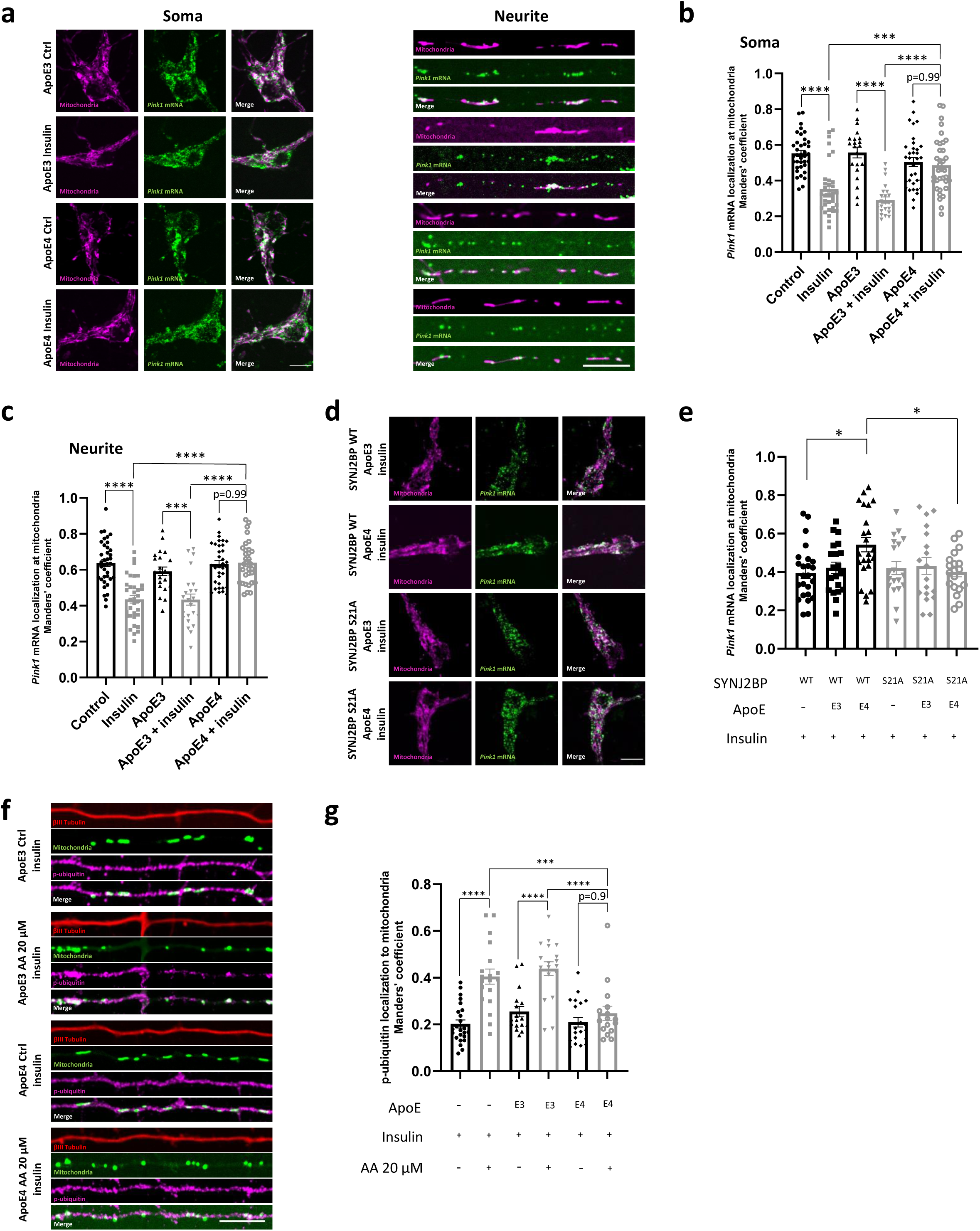
ApoE4 inhibits insulin-regulated *Pink1* mRNA localization and PINK1 activation. **a** Representative images of *Pink1* mRNA visualized by the MS2/PP7 split Venus method and mitoRaspberry with and without insulin (500 nM, 1 h) treament in the presence of ApoE3 (50 nM) and ApoE4 (50 nM) overnight, respectively, in the soma and neurites. **b** Quantification of the Manders’ colocalization coefficient for the overlap between the *Pink1* mRNA and mitochondrial channel in the soma as in **a**. One-way ANOVA followed by Tukey’s post hoc test; n = 21-30; p < 0.001 (***), p < 0.0001 (****). **c** Quantification as in **b** for neurites. One-way ANOVA followed by Tukey’s post hoc test; n = 22-37; p < 0.001 (***), p < 0.0001 (****). **d** Representative images of *Pink1* mRNA visualized by the MS2/PP7 split Venus method and mitoRaspberry upon overnight treatment with insulin as well as ApoE3 (50 nM) and ApoE4 (50 nM), respectively, combined with overexpression of SYNJ2BP WT and S21A, respectively. **e** Quantification of the Manders’ colocalization coefficient for the overlap between the *Pink1* mRNA and mitochondrial channel as in **d**. One-way ANOVA followed by Tukey’s post hoc test, n = 18-24, p < 0.05 (*). **f** Representative images of neurites overexpressing mito-meGFP cultured in the presence of insulin as well as ApoE3 (50 nM) and ApoE4 (50 nM) overnight, respectively, prior to treatment with 20 µM AA and stained with an antibody against phospho-ubiquitin (S65) and βIII tubulin. **g** Quantification of phospho-ubiquitin (S65) localization to mitochondria using the Manders’ colocalization coefficient as in **f**. One-way ANOVA followed by Tukey’s post hoc test; n = 17-24; p < 0.001 (***), p < 0.0001 (****). All data are expressed as mean±SEM. All data points represent single cells coming from ≥3 biological replicates. Scale bars, 10 µm.

## Discussion

Our study identifies metabolic control of *Pink1* mRNA localization in response to activation of IR signaling and downstream inhibition of AMPK. Upon insulin withdrawal, AMPK-mediated phosphorylation of the mitochondrial outer membrane protein SYNJ2BP favors its interaction with the RNA-binding protein SYNJ2 as well as the mitochondrial localization of the *Pink1* mRNA (Fig. 4 and 5). This tight association allows efficient transport of the transcript into neurites via mitochondrial hitch-hiking^5^. Fitting to its role as a master regulator of cellular metabolism, AMPK is known to impact several aspects of mitochondrial biology, ranging from activation of mitochondrial fission to suppression of retrograde transport and synaptic anchoring^51–53^. These mechanisms serve to increase mitochondrial presence in axons that undergo shortage of ATP due to synaptic activity. Our results predict that mitochondrial hitch-hiking will be increased under these conditions, while at the same time local PINK1 translation will be reduced (Fig. 6). This limits the amount of local mitophagy, which further serves to increase the local mitochondrial content, at the expense of retaining mildly damaged organelles (compare Fig. 6c,d). Local biogenesis of PINK1 in neurons is then activated by untethering of the *Pink1* transcript from mitochondria during times of glucose abundance, as signaled by the presence of insulin and inhibition of AMPK signaling (Fig. 1 and 2). During these times, neurons may generate ATP through glycolysis as well as via mitochondrial OXPHOS^54^, reducing the absolute necessity of mitochondria and opening a window for mitochondrial quality control via PINK1.

In our model, neuronal PINK1-dependent mitophagy would be regulated oppositely to bulk autophagy, which increases during fasting via activation of AMPK and inhibition of mTOR signaling^55^. This may be necessary in the morphologically complex environment of neurons, where mitostasis is particularly challenging^2^. Removal of a damaged organelle resident at a particular synapse will alter the local calcium dynamics^56^ and local mitophagy may therefore present a last resort that only will be activated if the mitochondrion is severely damaged. In line with this, PINK1/Parkin-mediated mitophagy is not observed under basal conditions in neurons^57, 58^, and while starvation reliably represses mTOR signaling in neurons, this does not lead to increased autophagy in cultured neurons^59^, in line with our observations. Inhibition of mTOR signaling also leads to repression of bulk translation, which is thought to decrease the energetic needs of the cell. Our study now adds an AMPK-dependent, parallel pathway that attenuates local PINK1 synthesis in neurons. Besides its role in mitophagy, PINK1 has also been described to positively stimulate the translation of other mitochondrially associated mRNAs encoding mitochondrial OXPHOS proteins^60^. Restriction of PINK1 biogenesis to times of energy abundance may in turn also synchronize the biogenesis of several mitochondrially associated transcripts and limit energy-consuming local translation during ATP-limiting conditions. Our results are in line with previous studies that connect PINK1-dependent mitophagy in neurons with the activity of the AKT kinase^61^. As this PI(3,4,5)P_3_ (PIP_3_)-activated kinase is a common downstream effector of several growth factor receptors, it is interesting to speculate that similar mechanisms also regulate *Pink1* mRNA localization upon signaling by other growth factors using this pathway. Intriguingly, PINK1 itself has been suggested to positively regulate the levels of PIP_3_, which promotes AKT activity^62^. This could build a positive feedback loop that ensures efficient AKT activity locally to inhibit AMPK and release more *Pink1* mRNA from its mitochondrial sequestration to amplify local PINK1 activity and mitophagy.

Importantly, we have seen that while low doses of AA required the presence of insulin signaling to fully activate mitophagy (Fig. 6c,d), strong mitochondrial impairment can still activate mitophagy even in the absence of insulin (Ext. Data Fig. 6a,b). The reduced levels of PINK1 protein upon insulin withdrawal detected by immunoblotting (Fig. 6e,f) may be sufficient to recruit and activate a small amount of Parkin (Fig. 6a,b). Parkin may then use non-phosphorylated ubiquitin to still demarcate the damaged mitochondrion, consistent with the observed lack of phospho-ubiquitin enrichment (Fig. 6g,h). It remains to be determined whether also other mitophagy pathways are actively upregulated to balance the reduced activity of the PINK1/Parkin-dependent pathway during energy-limiting conditions. One potential candidate is FUNDC1-mediated mitophagy. AMPK signaling activates the general autophagosome machinery including direct phosphorylation of the kinase ULK1 as part of the autophagy pre-initiation complex^63^. AMPK-phosphorylated ULK1 subsequently translocates to mitochondria and can phosphorylate and activate FUNDC1^64, 65^. This may represent one of the alternative delivery routes of mitochondria to the autophagolysosome under energy-limiting conditions. Also, Parkin has been shown to be phosphorylated by cytosolic ULK1^66^, which promotes the ability of PINK1 to activate Parkin. It remains to be determined whether ULK1-mediated phosphorylation of Parkin or FUNDC1 contributes to mitophagy in neurons.

This insulin-mediated regulation of *Pink1* mRNA tethering to mitochondria may also serve to resolve some of the controversial results in the field. Low doses of AA have been shown to be insufficient to elicit Parkin recruitment to mitochondria^67^ and, interestingly, the hippocampal mouse neurons used in this study were imaged in buffer not containing any growth factors. Also, in iPSC-derived neurons, differences in the efficacy of PINK1/Parkin-mediated mitophagy have been observed: While some studies were able to detect Parkin recruitment to mitochondria^68, 69^, others have argued that this pathway is non-functional in human iPSC-derived neurons^70^. Our results now suggest that differential media compositions may underlie some of the observed variability. It is interesting to speculate that insulin-triggered regulation of PINK1 activity *in vivo* also underlies the lack of PINK1/Parkin-dependent mitophagy observed in mice expressing a mitophagy reporter^57^. According to our model, basal neuronal PINK1 activation may be highest during the transient, post-prandial increase in blood insulin levels, and therefore potentially easy to miss in *ad libitum* fed mice.

Insulin resistance is frequently observed in AD pathology, and altered brain glucose metabolism often precedes the clinical onset of the disease^71^, yet how it is connected to the pathogenesis of the disease remains to be established. Likewise, AD pathology includes mitochondrial dysfunction, yet how mitochondrial dysfunction arises and how it is connected to AD risk-genes is unknown^72^. Our results now suggest that mitochondrial dysfunction in neurons may arise as a direct consequence of insulin resistance, at least *in vitro*. We used exogenous addition of the AD-risk factor ApoE4 to the cell culture medium to induce insulin resistance in cultured neurons, which successfully prevented the inhibition of AMPK upon insulin treatment, unlike addition of its homolog ApoE3 (Extended Data Fig. 7a). Accordingly, ApoE4 addition prevented the untethering of *Pink1* mRNA (Fig. 7a-c) and effective labelling of damaged mitochondria with phospho-ubiquitin in response to insulin (Fig. 7f,g), skewing the balance towards reduced PINK1 activity. Fittingly, mice transgenic for the human APOE ε4-allele display lower cleaved PINK1 levels in hippocampal neurons^73^, and total PINK1 as well as phospho-ubiquitin levels are reduced in human APOE ε4 carriers^74^, in line with our findings. Insufficient mitochondrial quality control may underlie the decreased mitochondrial respiration also observed in the cortex of aged ApoE4 but not ApoE3 mice^75^. However, as some of these changes may also occur via transcriptional regulation^76^, the proportion of PINK1 attenuation via AMPK-mediated sequestering of *Pink1* mRNA on mitochondria *in vivo* needs to be determined experimentally.

In general, our identification of a direct connection between insulin signaling and neuronal PINK1 biogenesis integrates into the ever-growing body of literature that connects failures in mitochondrial quality control and decline in insulin-related signaling pathways with general aging and age-related diseases^77^. AMPK activation extends the life span in multiple organisms including mice^78–81^, yet how AMPK activity can be used to prevent or treat neurodegenerative diseases remains controversial^82^. Our results reveal that both metabolic states (insulin signaling off/AMPK on vs. insulin signaling on/AMPK off) are important for proper quality control of neuronal mitochondria by the PINK1/Parkin-dependent pathway, suggesting that AMPK activation will have detrimental effects on mitochondrial quality control. While AMPK activation allows the efficient transport of *Pink1* mRNA to distal neurites, the second state then provides the cue to translate PINK1 and improve mitochondrial quality. Loss of either of these states, as seen pathologically during insulin resistance, will inevitably reduce the neuronal capacity to eliminate damaged organelles and contribute to the oxidative damage of the cell.

## Acknowledgments

We thank J. Lindner, C. Weiß and A. Schneider for technical support and the members of the Harbauer laboratory for their support and many fruitful discussions. We are grateful to R. Kasper/E. Laurell and M. Spitaler/M. Oster from the Imaging facilities of the MPIs for Biological Intelligence and Biochemistry, respectively, for assistance with live-cell imaging. We also thank S. Suppmann from the protein purification core and B. Steigenberger from the Mass Spectrometry facility for their assistance with protein purification and phospho-proteomics. ABH is supported by the Max Planck Society and the Deutsche Forschungsgemeinschaft (DFG, German Research Foundation, HA 7728/2-1 - ID 453679203) and under Germany’s Excellence Strategy within the framework of the Munich Cluster for Systems Neurology (EXC 2145 SyNergy – ID 390857198), as well as by the European Union (ERC StG Project 101077138 — MitoPIP).

## Author Contributions

ABH conceived of the project and wrote the manuscript together with JTH. JTH designed and conducted all experiments.

## Materials and Methods

### Mice

C57BL/6 mice were housed in the animal facility of the Max Planck Institute for Biological Intelligence, Martinsried. All mouse procedures were performed according to the regulation of the Government of upper Bavaria.

### Cell culture

#### Primary mouse neurons

Primary mouse cortical and hippocampal neurons were prepared as described^5^. After euthanizing the pregnant mouse with CO2, the E16.5 embryos were extracted from the abdomen. Cortices and hippocampi were dissected and placed in ice-cold dissociation medium (Ca^2+^-free Hank’s Balanced Salt Solution with 100 mM MgCl_2_, 10 mM kynurenic acid, and 100 mM HEPES). After enzymatic dissociation using papain/L-cysteine (Sigma-Aldrich), trypsin inhibitor (Abnova) was added to the tissue followed by trituration (10-15 times) with a p1000 pipet until the material dissipated into a single cell suspension. Neurons were resuspended in Neurobasal medium supplemented with B27 (Thermo Fisher Scientific), penicillin/streptomycin, and L-glutamine (NB+B27+PSG) and plated on 20 μg/mL poly-L-Lysine (PLL; Sigma-Aldrich) and 3.5 μg/mL laminin (Thermo Fisher Scientific) coated glass bottom plates (CellVis) or acid washed glass coverslips (1.5 mm, Marienfeld). 50 % of the medium was replaced every 5 days with fresh NB+B27+PSG. Transfections were performed at day *in vitro* (DIV) 5-7 and imaging at DIV7–DIV9.

#### Induced pluripotent stem cell (iPCS)-derived cortical neurons

Human iPSCs (cell line: HPSI0314i-hoik_1) were obtained from the Wellcome Trust Sanger Institute HipSci Repository. iPSCs were maintained in StemFlex medium (Thermo Fisher Scientific) on Matrigel (Corning)-coated plates. The cells were passaged at 80 % confluency using ReLeSR (Stem Cell Technologies) enzyme free passaging reagent.

iPSCs were differentiated into cortical neurons by overexpressing the transcription factor neurogenin-2 (NGN-2) as described^83^. In brief, iPSCs were dissociated into single cells using Accutase (Thermo Fisher Scientific) and plated at a density of 2.5*10^6^ per 10 cm dish (Falcon) in StemFlex supplemented with 10 µM Y-27632 (Tocris) on day −2. On day −1, iPSCs were transduced with lentivirus packaged pLV-TetO-hNGN2-eGFP-puro and FudeltaGW-rtTA plasmids in StemFlex medium. On day 0, the medium was replaced with N2/DMEM/F12/NEAA (N2 medium; Thermo Fisher Scientific) containing 10 ng/ml BDNF (PeproTech), 10 ng/ml NT-3 (PeproTech) and 0.2 µg/ml laminin (Thermo Fisher Scientific). Furthermore, 2 µg/ml doxycycline (Takara) was added on day 0 to induce TetO gene expression. On day 1, 1 µg/ml puromycin (Enzo Life Sciences) was added to the N2 medium in order to start a 24-hour puromycin selection period. On day 2, the medium was replaced with B27/Neurobasal-A/Glutamax (B27 medium; Thermo Fisher Scientific) containing 10 ng/ml BDNF, 10 ng/ml NT-3, 0.2 µg/ml laminin, 2 µg/ml doxycycline and 2 µM Ara-C (Sigma-Aldrich). From day 3 – 6, B27 medium containing 10 ng/ml BDNF, 10 ng/ml NT-3, 0.2 µg/ml laminin, 2 µg/ml doxycycline and 2 µM Ara-C was exchanged every other day. From day 7 onwards, cells were cultured in conditioned Südhof neuronal growth medium (NGN2 glial conditioned medium/B27/Neurobasal-A/Glucose/NaHCO3/L-Glutamine/Transferrin) containing 10 ng/ml BDNF, 10 ng/ml NT-3 and 0.2 µg/ml laminin. 50 % of the medium was exchanged every other day. On day 8, the iPSC-derived cortical neurons were dissociated using TrypLE Express (Thermo Fisher Scientific) for 5 min at 37 °C and re-plated in 20 μg/mL PLL (Sigma-Aldrich) and 3.5 μg/mL laminin (Thermo Fisher Scientific)-coated 6 well plates (Greiner) at a density of 2*10^6^ cells per well. 50 % of the medium was replaced every other day with fresh conditioned Südhof neuronal growth medium. All treatments were carried out on DIV14.

#### HEK293T cells

HEK293T cells were purchased from ATCC and cultured in Dulbecco’s modified Eagle medium (DMEM; Thermo Fisher Scientific) containing 10 % fetal bovine serum (FBS) in T75 flasks (Falcon). The cells were passaged at 80 % confluency using Trypsin (Thermo Fisher Scientific).

#### DNA constructs

Mito-mRaspberry-7, mito-meGFP, EBFP2-C1 (cell fill), YFP-Parkin, mito-mKeima, pPBbsr2-4031NES (AMPK FRET biosensor), pLV-TetO-hNGN2-eGFP-puro and FudeltaGW-rtTA were acquired from Addgene (55931, 172481, 54665, 23955, 131626, 105241, 79823 and 19780, respectively). Plasmids encoding shRNA against SYNJ2BP and AMPK α1 and α2 (TRCN0000139049, TRCN0000024000 and TRCN0000024046, respectively) as well as a control shRNA plasmid (TR30021) were purchased in pLKO from Sigma-Aldrich. PINK1-kinase dead-MS2-PP7, split-Venus and shRNA-resistant myc-tagged SYNJ2BP WT plasmids have previously been described^5^. SYNJ2BP mutations S21A and S21E were introduced by site-directed mutagenesis. The lentiviral packaging plasmids used for virus production pMDLg/pRRE, pRSV-Rev and pMD2.G were acquired from Addgene (12251, 12253 and 12259, respectively). For purification of SYNJ2BP from *E. Coli*, the cytosolic domain of rat SYNJ2BP WT (amino acids 1-110) was inserted into the bacterial expression vector pET19b and C-terminally tagged with 6xHis-tag. The SYNJ2BP S21A mutation was introduced by site-directed mutagenesis.

#### Protein purification of recombinant SYNJ2BP

Rosetta *E. coli* bacteria were transformed with the plasmid pET19b-SYNJ2BP-6xHis WT or S21A. Bacteria were inoculated in 15 ml ZY auto-induction medium overnight, before fermenter inoculation at OD 0.04. After 24 h, bacteria were harvested by centrifugation at 8000 rpm for 10 min at 4 °C to yield a pellet of 50 g. Half of the pellet was then resuspended in 100 ml lysis buffer, comprising of His-Binding Buffer (50 mM Na_3_PO_4_/Na_2_HPO_4_ pH 8, 500 mM NaCl, 10 mM imidazole, 10 % glycerol) supplemented with 1 mM AEBSF-HCl, 2 µg/ml Aprotinin, 1 µg/ml Leupeptin, 1 µg/ml Pepstatin, 2,4 U/ml Benzonase Corefa and 2 mM MgCl_2_. Bacteria were lysed by homogenization using an Avestin system and the lysate was cleared by centrifugation at 20500 rpm for 30 min at 4 °C.

The his-tagged proteins were purified via NiNTA-affinity chromatography using a linear gradient from 4 to 100 % elution buffer (50 mM Na_3_PO_4_/Na_2_HPO_4_ pH 8, 500 mM NaCl, 250 mM imidazole, 10 % glycerol) over 10 column volumes and collected in 1 ml fractions. Pooled fractions containing high amounts of recombinant SYNJ2BP were further purified by subsequent size-exclusion chromatography using a HiLoad 16/60 Superdex 75 column in SE/storage Buffer (20 mM Tris pH 7.2, 30 mM NaCl, 10 % glycerol). A final concentration step was carried out using Amicon Ultra 15 columns in two batches to a final concentration of 1 mg/ml and frozen at −80 °C.

#### *In vitro* phosphorylation assay with recombinant AMPK

For the *in vitro* phosphorylation assay, purified SYNJ2BP WT and SYNJBP S21A mutant protein were used. Prior to phosphorylation, recombinant SYNJ2BP (0.5 µg) was dephosphorylated using 1 µl of calf intestinal alkaline phosphatase (CIP) (NEB) in 1x CutSmart Buffer in a total volume of 10 µl. The reactions were incubated at 37 °C for 1 h and subsequently stopped by addition of 1x PhosStop (Merck). *In vitro* phosphorylation of dephosphorylated recombinant SYNJ2BP (0.5 µg in 10 µl) was performed in kinase reaction buffer (8 mM MOPS/NaOH pH 7, 200 µM EDTA) supplemented with 500 µM ATP (Serva), 200 µM AMP (Serva) and active recombinant AMPK (16 ng) (Sigma-Aldrich; 14-840) in a total volume of 30 µl. These samples were incubated at 30 °C for 2 h, shaking at 300 rpm. The reactions were stopped by addition of Laemmli sample buffer and boiling at 95 °C for 5 min.

#### *In vitro* phosphorylation assay with neuronal cytosolic extracts

Primary mouse cortical neurons were seeded in 6 well plates (Greiner) at a density of 2*10^6^ per well and maintained in NB+B27+PSG as described above. Prior to generation of cell lysates on DIV6, cells were either treated with 1 mM AICAR (Abcam) for 2 h, or were cultured for 2 h in medium lacking B27 (NB+PSG). After two washing steps with ice-cold PBS, neurons were lysed by dounce homogenization in a buffer containing 20 mM Tris/HCl pH 7.2, 30 mM NaCl, 10 mM MgCl_2_, 10 % glycerol, 1 mM EDTA, 200 µM PMSF and protease inhibitor cocktail (Roche) (80 µl/well). Lysates were cleared by centrifugation at 5000 rpm for 1 min at 4 °C. The supernatant (cytosolic extract) was collected and immediately used for *in vitro* phosphorylation assays or snap frozen and stored at −80 °C. For *in vitro* phosphorylation assays, 0.5 µg of purified and dephosphorylated SYNJ2BP (as described above) was mixed with 25 µl of cytosolic extract supplemented with 500 µM ATP in a final volume of 40 µl. After incubation at 30 °C and 300 rpm shaking for 2 h, reactions were stopped by addition of Laemmli sample buffer and boiling at 95 °C for 5 min.

#### Phos-Tag SDS-PAGE

*In vitro* phosphorylation assay samples of SYNJ2BP were analyzed using Phos-Tag sodium dodecyl sulfate polyacrylamide gel electrophoreses (Phos-Tag SDS-PAGE). To this end, standard discontinuous 20 % polyacrylamide gels were prepared with the modification that 50 µM Phos binding reagent acrylamide (PBR-A, APExBIO) and 100 µM Zn(NO_3_)_2_ were added to the separating gel. Electrophoresis was performed at 40 mA for 2 h. Prior to blotting, the gel was incubated for 30 min in standard transfer buffer containing 1 mM EDTA and for 10 min in transfer buffer without EDTA. For immunoblotting a PVDF membrane and a standard wet transfer protocol was used. Membranes were decorated with anti-SYNJ2BP rabbit antibody (1:500; Proteintech).

#### Immunoblotting for detection of PINK1 expression levels

Human iPSC-derived cortical neurons and HEK293T cells were seeded in 6 well plates (Greiner) at a density of 2*10^6^ cells and 0.5*10^6^ cells per well, respectively. iPSC-derived neurons were harvested on DIV14 and HEK293T cells 2 days after plating using lysis buffer containing 25 mM Tris/HCl pH 7.4, 150 mM NaCl, 1 % NP-40, 1 mM EDTA supplemented with 200 µM PMSF and protease inhibitor cocktail (Roche). Prior to lysis, iPSC-derived neurons were cultured for 2 h in medium lacking insulin or treated with 10 µM AKT inhibitor VIII (TCI) for 2 h. HEK293T cells were treated with either 1 µM GSK1904529A (Abcam) or 10 µM AKT inhibitor VIII (TCI) for 2 h. The samples were analysed by gel electrophoresis on a 12 % SDS-PAGE gel and immunoblotting using anti-PINK1 rabbit antibody (1:500; Novus Biologicals), anti-β-actin mouse (AC-74) antibody (1:500; Sigma-Aldrich) and anti-βIII-tubulin mouse (2G10) antibody (1:2000; Invitrogen).

#### Lentivirus production

Lentiviral particles were produced in HEK293T cells. On day −1, HEK293T cells were seeded in collagen (Sigma-Aldrich) coated-10 cm dishes (Falcon) at a density of 6*10^6^ cells per dish. After 24 h (day 0), cells were transfected with 5 µg of the packaging plasmid mix (pMDLg/pRRE, pRSV-Rev, pMD2.G; ratio 4:1:1) and 5 µg of the transfer plasmid (pLV-TetO-hNGN2-eGFP-puro, FudeltaGW-rtTA, myc-tagged SYNJ2BP WT, S21A or S21E) using TransIT-Lenti reagent (Mirus Bio). After 48 h (day 2), medium containing lentiviral particles was mixed with lentivirus precipitation solution (Alstem; ratio 4:1) and incubated at 4 °C overnight. On day 3, the lentiviral particles were collected by centrifugation at 1500xg for 30 min, resuspended in 1 ml ice-cold PBS per dish and stored at −80 °C.

#### Co-immunoprecipitation

Primary mouse cortical neurons were seeded in 6 well plates (Greiner) at a density of 2*10^6^ cells per well. On DIV1, neurons were transduced with lentivirus packaged myc-tagged SYNJ2BP WT plasmid. On DIV6, cells were cultured in insulin-free B27 medium for 2 hours and afterwards lysed (25 mM Tris/HCl pH 7.4, 150 mM NaCl, 1 % NP-40, 1 mM EDTA supplemented with 200 µM PMSF and protease inhibitor cocktail (Roche)) and cleared by centrifugation at 10000 rpm for 1 min. The supernatant was incubated either with or without CIP at 37 °C for 1 hour. Afterwards, the supernatant was incubated with 2 µg anti-SYNJ2BP antibody (proteintech)/ml lysate for 30 min at 4 °C. Protein A Sepharose beads (10 mg/sample) were blocked with 1 % BSA and added to the supernatant with the antibody. After 60 min incubation at 4 °C, beads were collected in columns and washed in a buffer containing 20 mM Tris/HCl pH 8, 140 mM NaCl, 5 mM MgCl_2_, 0.5 mM DTT, 0.1 % Triton-X and 200 µM PMSF. After five washing steps proteins were eluted by addition of Laemmli sample buffer and boiling at 95 °C for 2 min. The samples were analysed by gel electrophoresis on a 7.5 % −15 % gradient SDS-PAGE gel and immunoblotting using anti-SYNJ2BP rabbit antibody (1:500; Proteintech), anti-SYNJ2 rabbit antibody (1:500; Proteintech) and anti-βIII-tubulin mouse (2G10) antibody (1:2000; Invitrogen).

### Phospho-mass spectrometry

#### Sample Preparation

The cell pellets of mouse cortical neurons (10*10^6^ cells) lentivirally overexpressing myc-tagged SYNJ2BP WT were incubated with 200 µl of preheated SDC buffer (1 % SDC, 40 mM CAA, 10 mM TCEP in 100 mM Tris, pH 8.0). After incubation for 2 min at 95 °C, the samples were ultrasonicated for 10 min with 10 x 30 s at high intensity and 30 s pause between each cycle (Bioruptor Plus sonication system, Diogenode). Incubation and ultrasonication was repeated for a second time. Then, the sample was diluted 1:1 with MS grade water. Proteins were digested with 2 µg Lys-C for 4 h and then overnight at 37 °C with 4 µg trypsin. The solution of peptides was then acidified with TFA to a final concentration of 1 %, followed by desalting via Sep-Pak C18 5cc vacuum cartridges. The dried peptides were resuspended in 220 µl of equilibration buffer (80 % acetonitrile, 0.1 % TFA) shortly before the phospho-peptide enrichment, or were resuspended in buffer A (0.1 % FA) for the measurement of the total proteome.

#### Phospho-peptide enrichment

Phospho-peptide enrichment was performed in an automated fashion on an AssayMAP Bravo Platform (Agilent Technologies) using AssayMAP FE(III)-NTA cartridges (Agilent Technology). After priming the cartridges with 200 µl of priming buffer (100 % acetonitrile, 0.1 % TFA) and equilibration with 250 µl of equilibration buffer (80 % acetonitrile, 0.1 % TFA), the sample were loaded onto the cartriges followed by washing with 250 µl equilibration buffer and elution with 35 µl of elution buffer (10 % NH_3_ in water). Eluted peptides were acidified with 35 µl of 10 % FA. Samples were concentrated in a SpeedVac for 45 min at 37 °C.

#### LC MS/MS data acquisition

Peptides were loaded onto a 30-cm column (inner diameter: 75 µm; packed in-house with ReproSil-Pur C18-AQ 1.9-micron beads, Dr. Maisch GmbH) via the autosampler of the Thermo Easy-nLC 1200 (Thermo Fisher Scientific) at 60 °C. Using the nanoelectrospray interface, eluting peptides were directly sprayed onto the benchtop Orbitrap mass spectrometer Exploris 480 (Thermo Fisher Scientific). Peptides were loaded in buffer A (0.1 % FA) and separated at a flow rate of 300 nL/min using an increasing gradient of buffer B (80 % acetonitrile, 0.1 % FA) from 5 % to 30 % over 105 min followed by an increase to 65 % over 5 min then 95 % over the next 5 min. Finally, percentage of buffer B was maintained at 95 % for another 5 min. The mass spectrometer was operated in a data-dependent mode with a survey scans from 300 to 1650 m/z (resolution of 60000 at m/z =200), and up to 15 of the top precursors were selected and fragmented using higher energy collisional dissociation (HCD with a normalized collision energy of value of 28). The MS2 spectra were recorded at a resolution of 30000 (at m/z = 200). AGC target for MS and MS2 scans were set to 3E6 and 1E5 respectively within a maximum injection time of 25 ms for MS1. Injection time for the MS2 scans was set to “Auto”. Dynamic exclusion was set to 30 ms.

#### Data analysis

Raw data were processed using the MaxQuant computational platform (version 2.0.1.0)^84^ with standard settings applied. Shortly, the peak list was searched against the reviewed mouse proteome data base and the sequence of SYNJ2BP (rat) with an allowed precursor mass deviation of 4.5 ppm and an allowed fragment mass deviation of 20 ppm. MaxQuant by default enables individual peptide mass tolerances, which was used in the search. Cysteine carbamidomethylation was set as static modification. Methionine oxidation, N-terminal acetylation, deamidation on asparagine and glutamine and phosphorylation on serine, threonine and tyrosine were set as variable modifications. The match-between runs option was enabled. The LFQ (label free quantification) algorithm was used for quantification of proteins and the iBAQ algorithm was used for calculation of approximate abundances for the identified proteins.

#### RNA isolation and RT-qPCR

Primary mouse cortical neurons were seeded in 6 well plates (Greiner) at a density of 2*10^6^ per well and maintained in NB+B27+PSG as described above. On DIV6, neurons were treated with or without 20 µM CC (Abcam) for 2 h prior to isolation of RNA using the QIAGEN RNeasy Mini Kit. Complementary DNA (cDNA) was generated using the qScript^TM^ cDNA SuperMix (Quantabio) followed by a RT-qPCR assay using the PerfeCTa SYBR® Green FastMix (Quantabio) in a Mic (magnetic induction cycler) PCR machine (Bio Molecular Systems). *Pink1* transcript levels upon CC treatment were determined relative to β*-actin* and control treatment using the comparative Ct method (formula: 2^-ΔΔCt^).

#### Live cell mRNA imaging

Live cell mRNA imaging in primary mouse hippocampal neurons was performed as described^5^. Briefly, primary neurons were seeded in 24-well glass bottom plates (CellVis) at a density of 100*10^3^ cells per well and maintained as described above. On DIV5-7, neurons were transfected for 20 min using lipofectamine 2000 transfection reagent (Thermo Fisher Scientific) in NB+PSG. The optimal ratio between the Pink1 construct containing the MS2 and PP7 binding sites and the split-Venus construct containing the respective coat proteins was empirically determined to be 4:1. Furthermore, 4 µg of the Pink1 construct was transfected to achieve export from the nucleus. Constructs were expressed for 2 days, except in the case of cells co-transfected with SYNJ2BP or AMPK shRNAs, which were imaged after 4 days to provide enough time for effective knock down of the respective proteins. On DIV9, imaging was performed in Hibernate E medium (BrainBits) at the Imaging Facility of the Max Planck Institute for Biological Intelligence, Martinsried, with an Eclipse Ti2 spinning disk microscope (Nikon) equipped with a DS-Qi2 high-sensitivity monochrome camera (Nikon) using a 60×/NA 1.2 oil immersion lens and NIS-Elements software (Nikon). Prior to imaging, respective treatments of the neurons were performed: 20 µM CC (Abcam) for 2 h, 1 mM AICAR (Abcam) for 2 h, 500 nM insulin (Sigma-Aldrich) for 1 h, 1 µM GSK1904529A (Abcam) for 2 h, 1 µM Wortmannin (EMD Millipore) for 2 h, 10 µM AKT inhibitor VIII (TCI) for 2 h, 50 nM ApoE3 (PeproTech) overnight and 50 nM ApoE4 (PeproTech) overnight.

#### Proximity Ligation Assay

The proximity ligation assay was performed according to the manufacturer’s instructions (Sigma-Aldrich) and as described^5^. Briefly, primary mouse hippocampal neurons were seeded on glass coverslips (1.5 mm, Marienfeld) at a density of 100*10^3^ and maintained as described above. Prior to fixation on DIV7, respective treatments of the neurons were performed: 20 µM CC (Abcam) or 1 mM AICAR (Abcam) were added to the cells cultured in either NB+B27+PSG or Hibernate E medium (BrainBits) for 2 h. 500 nM insulin (Sigma-Aldrich) and 1 µM GSK1904529A (Abcam) were added to NB+27+PSG for 1 h and 2 h, respectively. For starvation (minus insulin), neurons were cultured for 2 h in NB+B27+PSG lacking insulin. Furthermore, neurons were incubated with 100 nM MitoTracker Deep Red (Invitrogen) for 20 min at 37 °C followed by three washes in the respective medium. Afterwards, neurons were fixed with 4 % paraformaldehyde for 15 min and permeabilized with 0.3 % Triton X-100/PBS for 10 min followed by a 1 h incubation with Duolink blocking solution at 37 °C in a humidity chamber. Neurons were incubated with primary antibodies (anti-SYNJ2BP mouse antibody, 1:50, Sigma-Aldrich; anti-SYNJ2 rabbit antibody, 1:50, Proteintech) diluted in Duolink antibody diluent at 4 °C overnight. On the next day, the neurons were washed two times with Buffer A (0.01 M Tris, 0.15 M NaCl and 0.05 % Tween 20) at RT for 5 min before incubation with the Duolink PLA Probes (anti-rabbit plus and anti-mouse minus) at 37 °C for 1 h. After two washes with Buffer A at RT for 5 min, the neurons were incubated with Duolink ligation solution at 37 °C for 30 min. This was followed by two more washes with Buffer A at RT for 5 min before incubation with Duolink amplification solution at 37 °C for 100 min. After two washes with Buffer B (0.2 M Tris, 0.1 M NaCl) at RT for 10 min and a final wash with 0.01x Buffer B for 1 min the coverslips were mounted in Fluoromount G (Thermo Fisher Scientific) and imaged at the Imaging Facility of the Max Planck Institute for Biological Intelligence, Martinsried, with a Nikon Ti2 spinning disk microscope using a 60x/NA 1.40 oil immersion objective. The number of PLA puncta per soma was quantified.

#### Fluorescence Lifetime Imaging (FLIM)

Primary mouse hippocampal neurons were seeded in 24-well glass bottom plates (CellVis) at a density of 100*10^3^ cells per well and maintained as described above. On DIV6, neurons were transfected for 20 min with the AMPK FRET biosensor pPBbsr2-4031NES^42^ using lipofectamine 2000 transfection reagent (Thermo Fisher Scientific) in NB+PSG. On the day before imaging, neuronal medium was switched to B27 medium supplemented either with or without insulin as well as with or without 50 nM ApoE3 (PeproTech) or 50 nM ApoE4 (PeproTech). On DIV8, imaging was performed in Hibernate E medium (BrainBits) supplemented with or without 500 nM insulin as well as with or without ApoE3 or ApoE4 at the Imaging Facility of the Max Planck Institute of Biochemistry, Martinsried, on a LEICA (Wetzlar, Germany) SP8 FALCON confocal laser scanning microscope equipped with an HCX PL APO 63x/1.2 motCORR CS water immersion objective using the LAS-X software (version 3.5.5). 2 h prior to imaging, neurons were treated with the different inhibitors of the insulin signaling cascade: 1 µM GSK1904529A (Abcam), 1 µM Wortmannin (EMD Millipore) and 10 µM AKT inhibitor VIII (TCI). To determine AMPK activity, the donor (super enhanced cyan fluorescent protein) fluorescence lifetime of the AMPK FRET biosensor pPBbsr2-4031NES was measured by FLIM at 37 °C. Neurons were excited with a pulsed diode laser (PicoQuant) at 440 nm and photon arrival times of a maximum of 1000 photons per brightest pixel were detected between 470 and 512 nm. The phasor analysis approach was used to determine the fluorescence lifetime of the donor^85, 86^.

### Mitophagy assays

#### Phospho-ubiquitin stainings

Primary mouse hippocampal neurons were seeded on glass coverslips (1.5 mm, Marienfeld) at a density of 50*10^3^ and maintained as described above. On DIV5-7, neurons were transfected for 20 min with mitochondrially-targeted meGFP as well as SYNJ2BP WT, S21A or S21E in case of the rescue experiments using lipofectamine 2000 transfection reagent (Thermo Fisher Scientific) in NB+PSG. Constructs were expressed for 2 days. On the day before fixation, neuronal medium was switched to B27 medium supplemented with or without insulin. The next day, neurons were pre-treated with or without 10 nM Torin-2 (Sigma-Aldrich) or 1 mM AICAR (Abcam) for 30 min and 2 h, respectively, prior to addition of 20 µM AA (Alfa Aesar) or ethanol for 45 min. Neurons were fixed with 4 % paraformaldehyde for 15 min, permeabilized with 0.3 % Triton X-100/PBS for 10 min and then incubated in blocking buffer (1 % BSA/PBS) for 1 h at RT. Afterwards, neurons were incubated with primary antibodies diluted in blocking buffer at 4 °C overnight. Primary antibodies and their dilutions were as follows: anti-phospho-ubiquitin S65 rabbit antibody (1:200; Millipore) and anti-βIII tubulin mouse (2G10) antibody (1:1000; Invitrogen). After three washes in PBS at RT for 5 min each, neurons were incubated with secondary fluorescent antibodies (Alexa Fluor 568 and 647) diluted in blocking buffer for 2 h at RT. After three washes in PBS at RT for 5 min each, the coverslips were mounted in Fluoromount G (Invitrogen) and imaged at an Eclipse Ti2 spinning disk microscope (Nikon) using a 60x/NA 1.40 oil immersion objective.

#### Parkin translocation

Primary mouse hippocampal neurons were seeded in 24-well glass bottom plates (CellVis) at a density of 100*10^3^ cells per well and maintained as described above. On DIV7, neurons were transfected for 20 min with mitochondrially-targeted mRaspberry, a cytosolic BFP (EBFP2-C1) and YFP-Parkin using lipofectamine 2000 transfection reagent (Thermo Fisher Scientific) in NB+PSG. Constructs were expressed for 2 days. On the day before imaging, neuronal medium was switched to B27 medium supplemented either with or without insulin. On DIV9, imaging was performed in Hibernate E medium (BrainBits) supplemented with or without 500 nM insulin at the Imaging Facility of the Max Planck Institute for Biological Intelligence, Martinsried, with an Eclipse Ti2 spinning disk microscope (Nikon) equipped with a DS-Qi2 high-sensitivity monochrome camera (Nikon) using a 60×/NA 1.2 oil immersion lens and NIS-Elements software (Nikon). Images of the same neurons were taken before and 30 min after addition of 20 µM AA (Alfa Aesar).

#### Mito-mKeima imaging

Primary mouse hippocampal neurons were seeded in 24-well glass bottom plates (CellVis) at a density of 100*10^3^ cells per well and maintained as described above. On DIV6, neurons were transfected for 20 min with mitochondrially-targeted mKeima using lipofectamine 2000 transfection reagent (Thermo Fisher Scientific) in NB+PSG. Constructs were expressed for 2 days. On the day before imaging, neuronal medium was switched to B27 medium supplemented with or without insulin. On DIV8, imaging was performed in Hibernate E medium (BrainBits) supplemented with or without 500 nM insulin at the Imaging Facility of the Max Planck Institute of Biochemistry, Martinsried, with a LEICA (Wetzlar, Germany) SP8 FALCON confocal laser scanning microscope equipped with an HCX PL APO 63x/1.2 motCORR CS water immersion objective using the LAS-X software (version 3.5.5). 45 min before imaging, AA (Alfa Aesar; 5 nM or 20 µM) or ethanol was added to the neurons. mKeima green was excited at 442 nm and mKeima red at 550 nm. The emission was detected sequentially from 555-620 nm for both excitation wavelengths.

#### Quantification and statistical analysis

Data are expressed as mean ± SEM. Statistical analysis was performed in GraphPad Prism version 9.1.0 for Windows, GraphPad Software, San Diego, California USA, www.graphpad.com. A student’s t-test was used for comparison of two conditions, while for multiple conditions a one-way ANOVA followed by Tukey’s or Dunnett’s multiple comparisons was performed. P<0.05 was considered significant (*), with further levels defined as p<0.01 (**), p<0.001 (***) and p<0.0001 (****). For statistically non-significant comparisons, p-values are given in the figure. Western Blot images were acquired using the Invitrogen iBright FL1000 Imaging System (Thermo Fisher Scientific) and quantified in Fiji/Image (National Institutes of Health)^87^.

Quantification of microscopy data was performed in Fiji/ImageJ. For the *Pink1* mRNA imaging, the Manders’ colocalization coefficient of the Venus and mitochondrial channel was analyzed in z-stack images using the ‘Just Another Colocalization Plugin’(JaCoP)^88^ as described^5, 89^. For neurites, images were straightened with a 20 pixel margin after maximum z-projection. For cell bodies, a square-shaped region including the entire cell body was chosen and no maximum z-projection was performed. For the rotated Venus quantification, a 10 by 10 µm square was chosen within the cell body based on the mitochondrial signal to exclude the nucleus. The Manders’ colocalization coefficient was quantified with and without rotation of the Venus channel. Manders’ coefficients were exported to Excel and plotted in GraphPad Prism.

P-ubiquitin stainings were analyzed using Fiji/ImageJ. The Manders’ colocalization coefficient of the p-ubiquitin and mitochondrial signal was analyzed using the JaCoP plugin. The images of intact neurites based on the βIII-tubulin signal were straightened with a 20 pixel margin. The Manders’ colocalization coefficients were exported to Excel and plotted in GraphPad Prism.

Parkin images were analyzed in Fiji/ImageJ. The neurite images were straightened with a 20 pixel margin. To determine Parkin-positive mitochondria before and after AA treatment, plots of fluorescence intensity versus the position along neurites were generated for both the mitochondrial and the Parkin channels using the plot profile function of Fiji/ImageJ. A mitochondrion was considered Parkin positive when the intensity of the respective YFP-Parkin peak was at least twice as high as the baseline intensity level.

Mito-mKeima images were analyzed using Fiji/ImageJ. A square-shaped region including the entire cell body was chosen. After creating a thresholded image, mitochondria were identified using the particle analyzer. The intensities of the non-thresholded images were calculated and the mean gray values were exported to Excel. The mitophagy index was determined in Excel by calculating the ratio of the mean gray values: [mKeima red/(mKeima green + mKeima red)].

---

**Extended Data Fig. 1 corresponding to Fig. 1.**
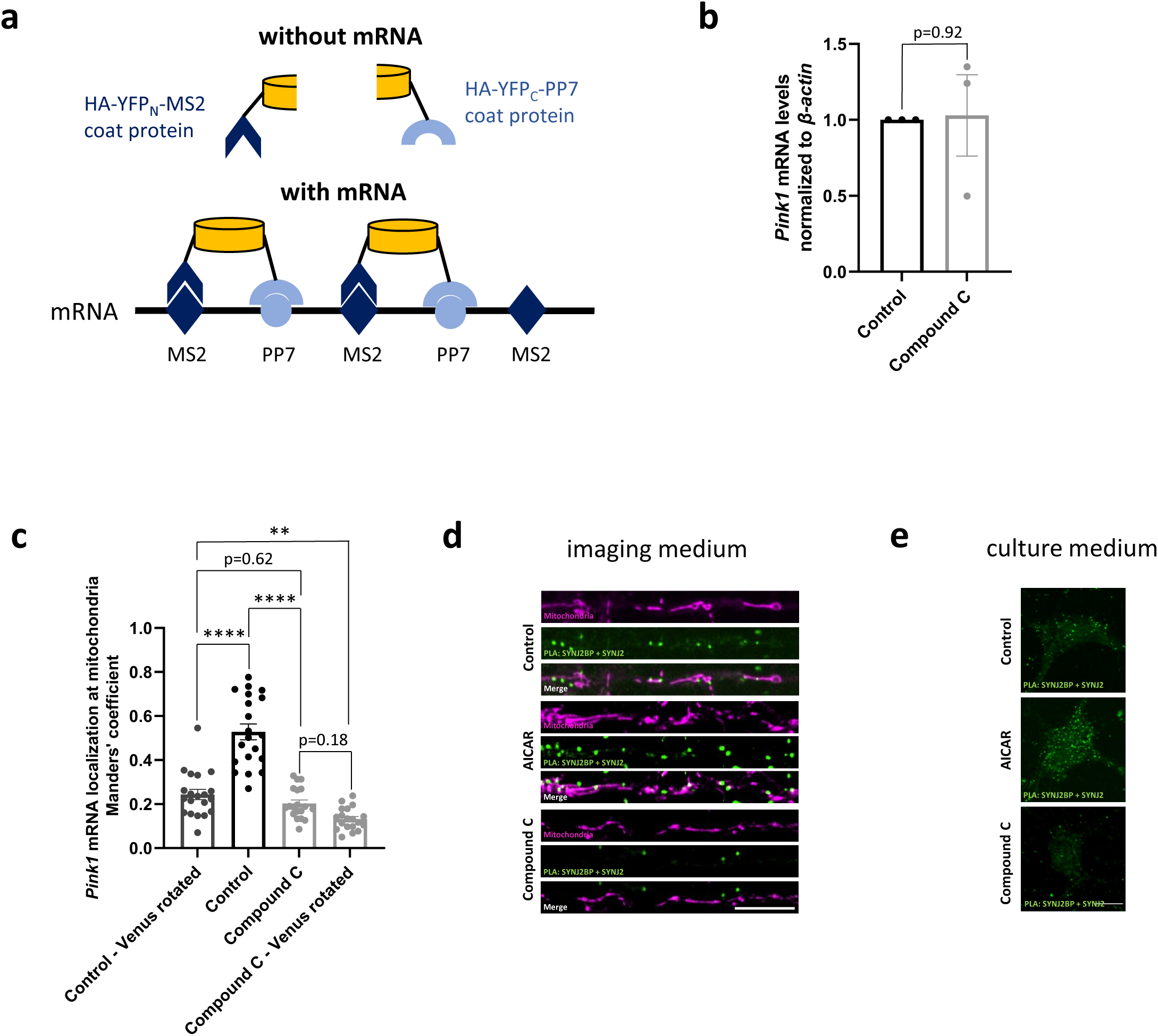
AMPK signaling regulates *Pink1* mRNA localization to mitochondria. **a** Schematic of the MS2/PP7-split Venus method for mRNA imaging. **b** RT-qPCR of *Pink1* transcript levels normalized to β-*actin* from primary cortical neurons treated with or without 20 µM CC for 2 h. Two-tailed student’s t-test; n = 3. **c** Live-cell imaging of *Pink1* mRNA localization in hippocampal neurons using the MS2/PP7-split Venus method. Quantification of the Manders’ colocalization coefficient for the overlap between the *Pink1* mRNA and the mitochondrial channel upon CC (20 µM, 2 h) treatment is shown for the soma. The analysis has been performed on a 10 by 10 µm square in the soma. ‘Venus rotated’ indicates that the *Pink1* mRNA channel had been rotated 90 degrees before quantification. One-way ANOVA followed by Tukey’s post hoc test; n = 19-20; p < 0.01 (**), p < 0.0001 (****). **d** Representative images of neurites displaying the PLA and mitotracker signal upon control, AICAR (1 mM, 2 h) and CC (20 µM, 2 h) treatment in imaging medium (Hibernate E). **e** Representative images of neuronal somas displaying the PLA signal upon control, AICAR (1 mM, 2 h) and CC (20 µM, 2 h) treatment in full culture medium. All data are expressed as mean±SEM. All data points represent single cells coming from ≥3 biological replicates. Scale bars, 10 µm.

**Extended Data Fig. 2 corresponding to Fig. 2.**
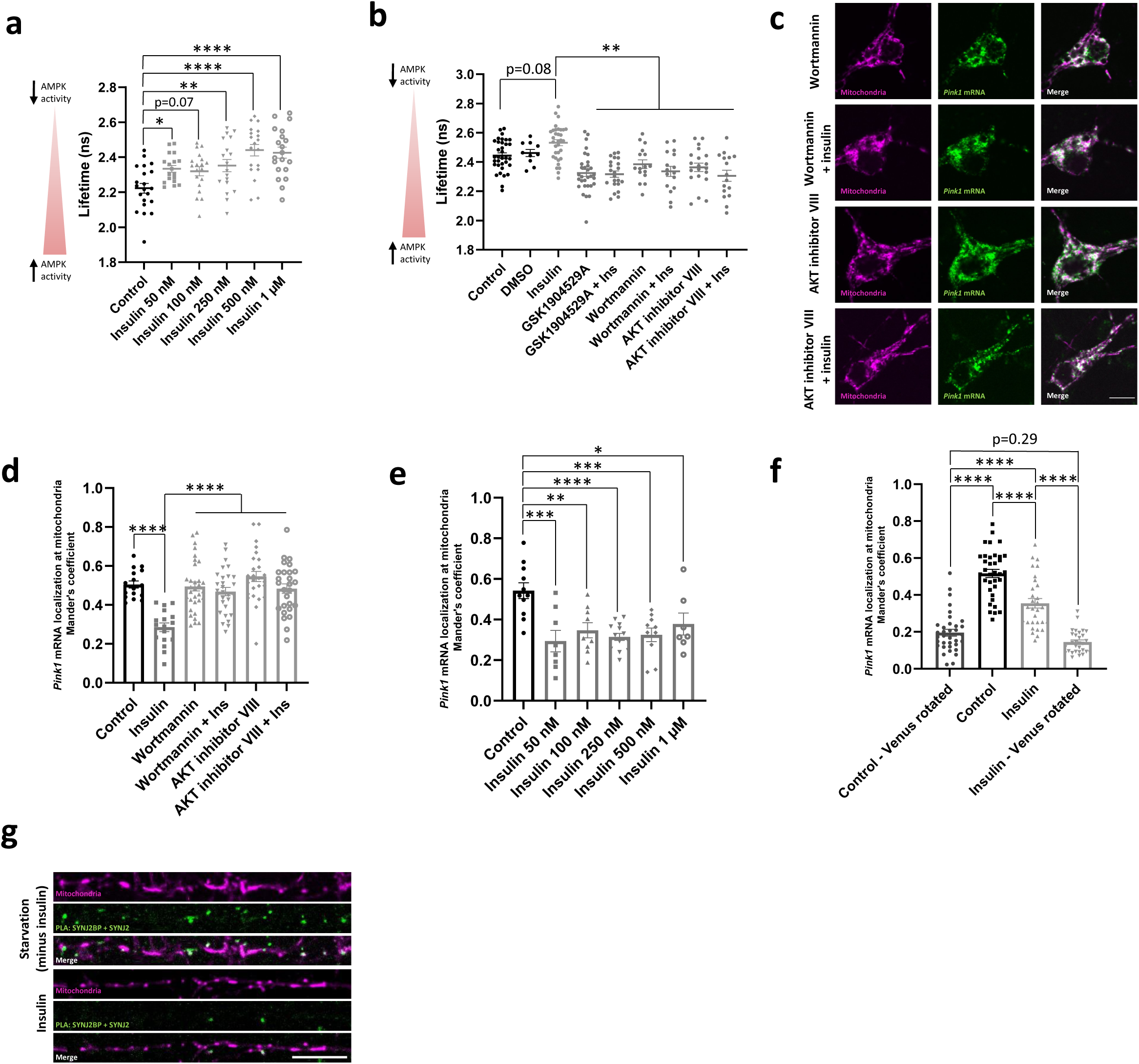
Insulin signaling inhibits AMPK and regulates *Pink1* mRNA localization to mitochondria. **a** Quantification of fluorescent lifetime imaging of the FRET-based AMPK activity sensor in neurons treated with increasing insulin concentrations for 1 h. One-way ANOVA followed by Dunnett’s post hoc test; n = 18-21; p < 0.05 (*), p < 0.01 (**), p < 0.0001 (****). **b** Quantification of fluorescent lifetime imaging of the FRET-based AMPK activity sensor in neurons treated with insulin in presence or absence of inhibitors of the insulin signaling pathway: IR inhibitor GSK1904529A (1 µM, 2 h), PI3K inhibitor Wortmannin (1 µM, 2 h) and AKT inhibitor VIII (10 µM, 2 h). One-way ANOVA followed by Tukey’s post hoc test; n = 10-34; p < 0.01 (**). **c** Representative images of *Pink1* mRNA visualized by the MS2/PP7 split Venus method and mitoRaspberry upon insulin (500 nM, 1 h) addition with or without pre-treatment with the PI3K inhibitor Wortmannin (1 µM, 2 h) and the AKT inhibitor VIII (10 µM, 2 h) in the soma. **d** Quantification of the Manders’ colocalization coefficient for the overlap between the *Pink1* mRNA and mitochondrial channel in the soma. One-way ANOVA followed by Tukey’s post hoc test; n = 16-32; p < 0.0001 (****). **e** Quantification of the Manders’ colocalization coefficient for the overlap between the *Pink1* mRNA and mitochondrial channel in neurons treated with increasing insulin concentrations for 1 h. One-way ANOVA followed by Dunnett’s post hoc test; n = 8-14; p < 0.05 (*), p < 0.01 (**), p < 0.001 (***), p < 0.0001 (****). **f** Live-cell imaging of *Pink1* mRNA localization in hippocampal neurons using the MS2/PP7-split Venus method. Quantification of the Manders’ colocalization coefficient for the overlap between the *Pink1* mRNA and the mitochondrial channel upon insulin (500 nM, 1 h) treatment is shown for the soma. The analysis has been performed on a 10 by 10 µm square in the soma. ‘Venus rotated’ indicates that the *Pink1* mRNA channel had been rotated 90 degrees before quantification. One-way ANOVA followed by Tukey’s post hoc test; n = 30-35, p < 0.0001 (****). **g** Representative images of neurites displaying the PLA and mitotracker signal upon starvation (minus insulin, 2 h) and insulin (500 nM, 1 h) treatment. All data are expressed as mean±SEM. All data points represent single cells coming from ≥3 biological replicates. Scale bars, 10 µm.

**Extended Data Fig. 3 corresponding to Fig. 3.**
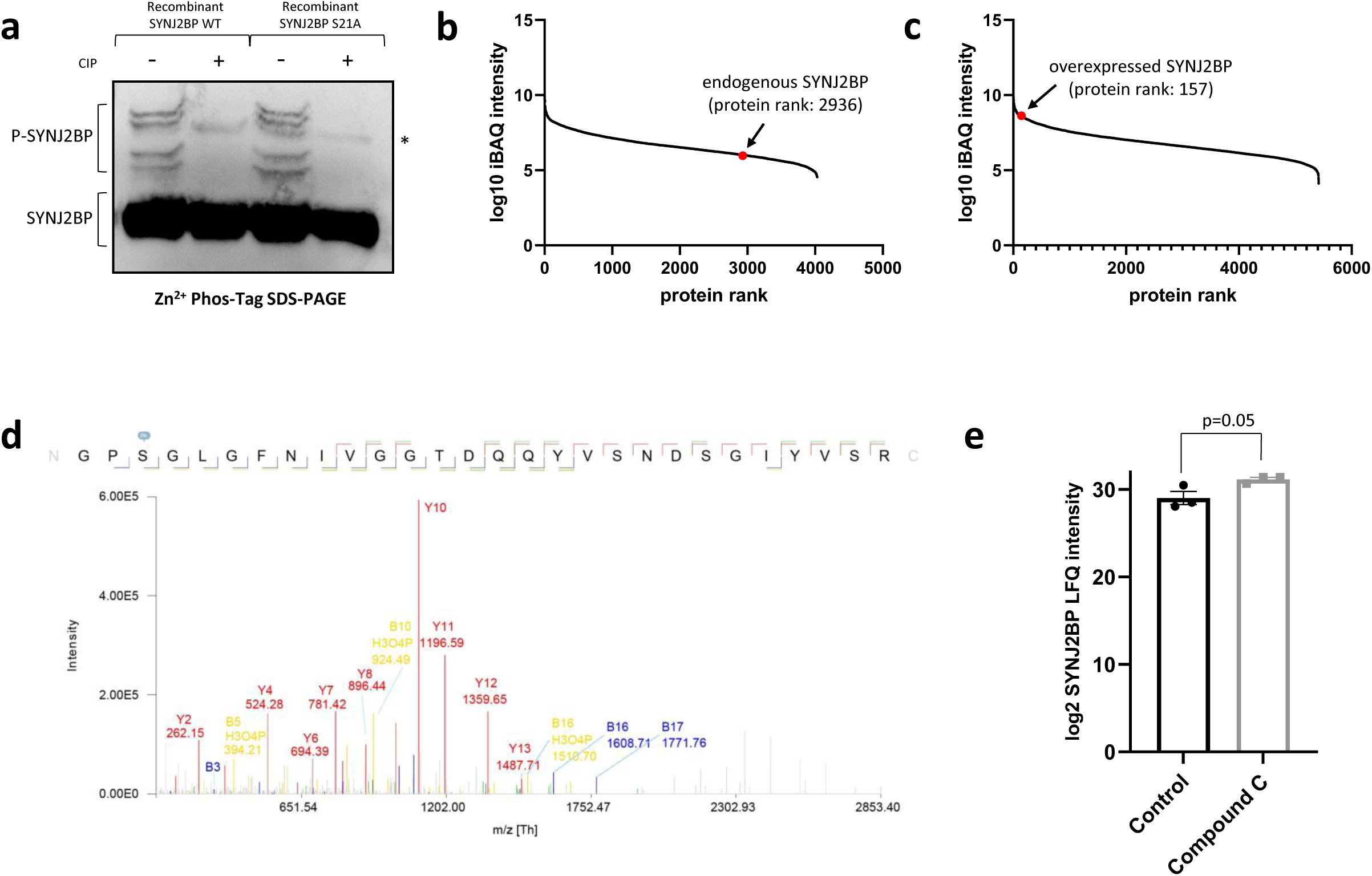
AMPK phosphorylates SYNJ2BP in its PDZ domain. **a** Dephosphorylation of recombinant SYNJ2BP WT and S21A, respectively, using CIP analyzed on a Zn^2+^-Phos-Tag SDS-PAGE and decorated with a SYNJ2BP antibody. Note the appearance of slower migrating species that disappear upon addition of CIP indicating phosphorylated forms of SYNJ2BP. The asterisk (*) denotes an unspecific band that is not responsive to CIP treatment. **b-c** The number of quantified proteins upon LC MS/MS analysis is plotted against the log10-transformed intensity-based absolute quantification (iBAQ) values. The iBAQ values represent an estimate of the molar abundance of proteins within the sample. Note, the protein rank of endogenous SYNJ2BP in cortical neurons (**b**) is relatively low, which can be improved by lentiviral overexpression of myc-tagged SYNJ2BP WT (**c**). **d** Annotated MS/MS spectrum of the peptide GPSGLGFNIVGGTDQQYVSNDSGIYVSR of lentivirally overexpressed myc-tagged SYNJ2BP in cortical neurons. Note, ion B3 represents the ion containing the phosphorylation at S21 of SYNJ2BP. **e** Log2-transformed LFQ intensities of SYNJ2BP upon LC MS/MS analysis in primary cortical neurons that lentivirally overexpress myc-tagged SYNJ2BP WT and are cultured in insulin-free medium and are treated with or without the AMPK inhibitor CC (20 µM, 2 h). Two-tailed student’s t-test; n = 3. All data are expressed as mean±SEM. All data points represent biological replicates.

**Extended Data Fig. 4 corresponding to Fig. 4.**
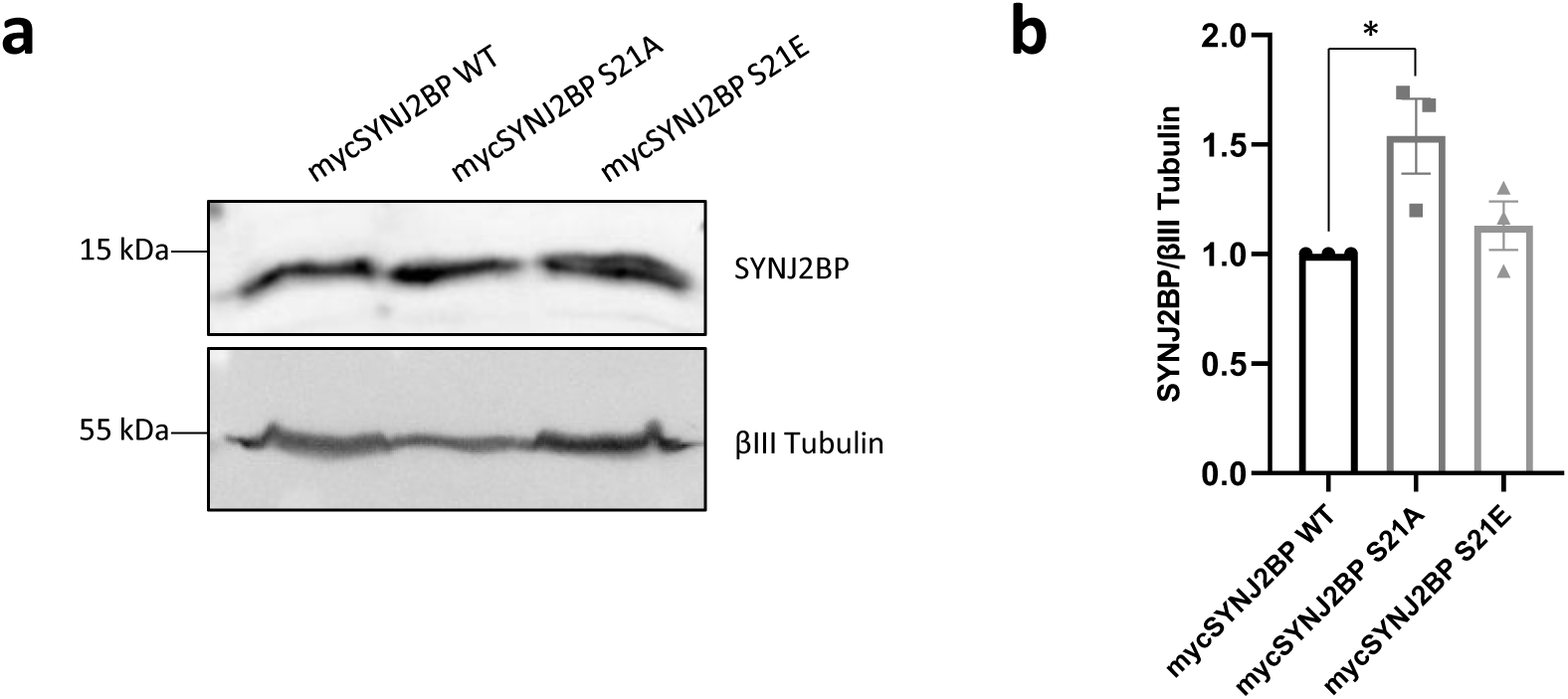
Expression levels of myc-tagged SYNJ2BP WT, S21A and S21E constructs. **a** Representative immunoblot images of cortical neurons that lentivirally overexpress myc-tagged SYNJ2BP WT, S21A and S21E, respectively. **b** Quantification of the SYNJ2BP protein bands normalized to the respective βIII tubulin bands. One-way ANOVA followed by Tukey’s post hoc test; n = 3; p < 0.05 (*). All data are expressed as mean±SEM. All data points represent biological replicates.

**Extended Data Fig. 5 corresponding to Fig. 5.**
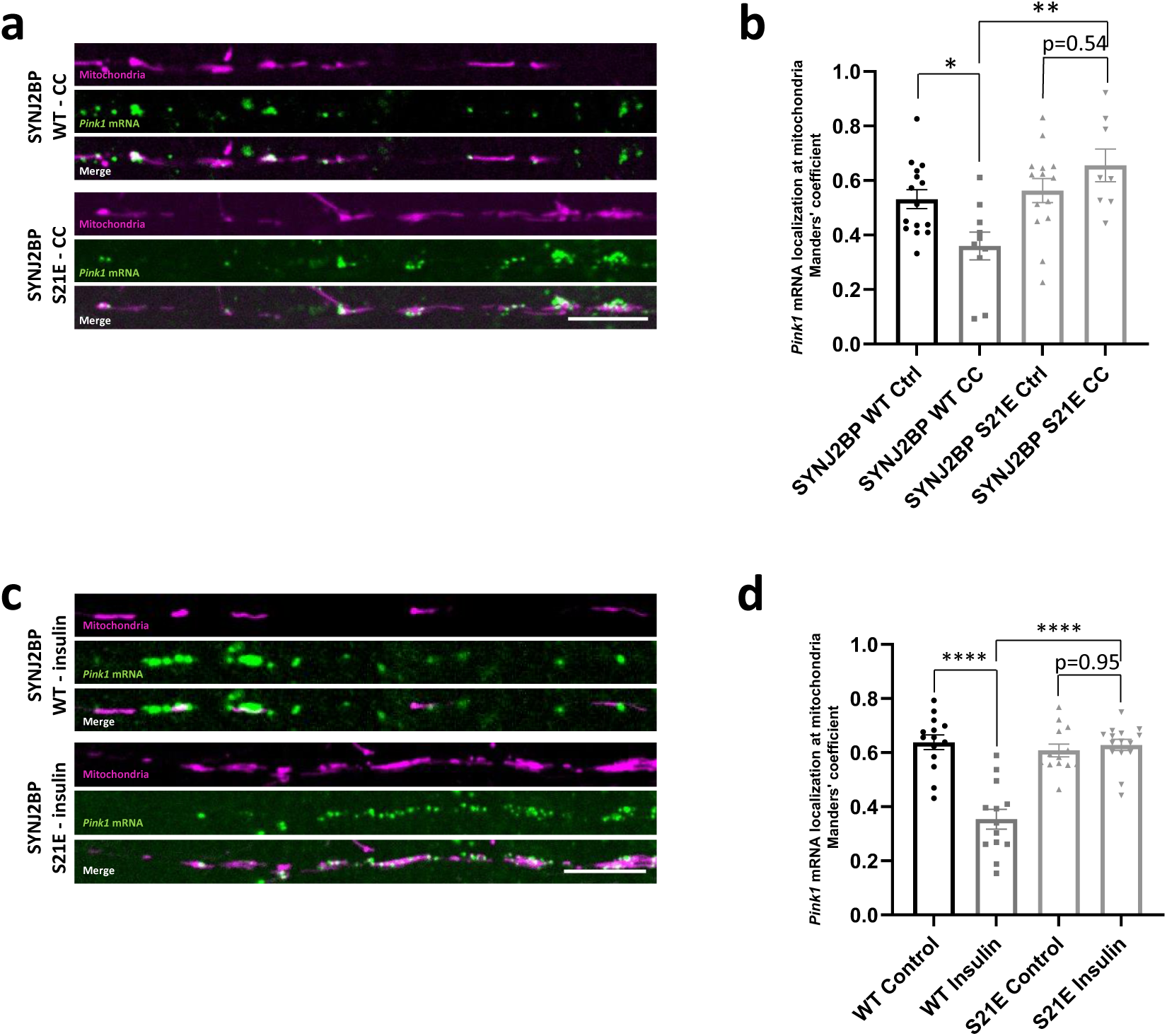
Phospho-mimetic SYNJ2BP restores mitochondrial *Pink1* mRNA localization upon AMPK inhibition. **a** Representative images of *Pink1* mRNA visualized by the MS2/PP7 split Venus method and mitoRaspberry in neurites upon CC (20 µM, 2 h) treatment combined with overexpression of SYNJ2BP WT and S21E, respectively. **b** Quantification of the Manders’ colocalization coefficient for the overlap between the *Pink1* mRNA and mitochondrial channel in neurites overexpressing SYNJ2BP WT or S21E and treated with or without CC (20 µM, 2 h). One-way ANOVA followed by Tukey’s post hoc test; n = 8-15; p < 0.05 (*), p < 0.01 (**). **c** Representative images of *Pink1* mRNA and mitoRaspberry in neurites upon insulin (500 nM, 1 h) treatment combined with overexpression of SYNJ2BP WT and S21E, respectively. **d** Quantification of the Manders’ colocalization coefficient for the overlap between the *Pink1* mRNA and mitochondrial channel in neurites overexpressing SYNJ2BP WT or S21E and treated with or without insulin (500 nM, 1 h). One-way ANOVA followed by Tukey’s post hoc test; n = 13-15; p < 0.0001 (****). All data are expressed as mean±SEM. All data points represent single cells coming from ≥3 biological replicates. Scale bars, 10 µm.

**Extended Data Fig. 6 corresponding to Fig. 6:**
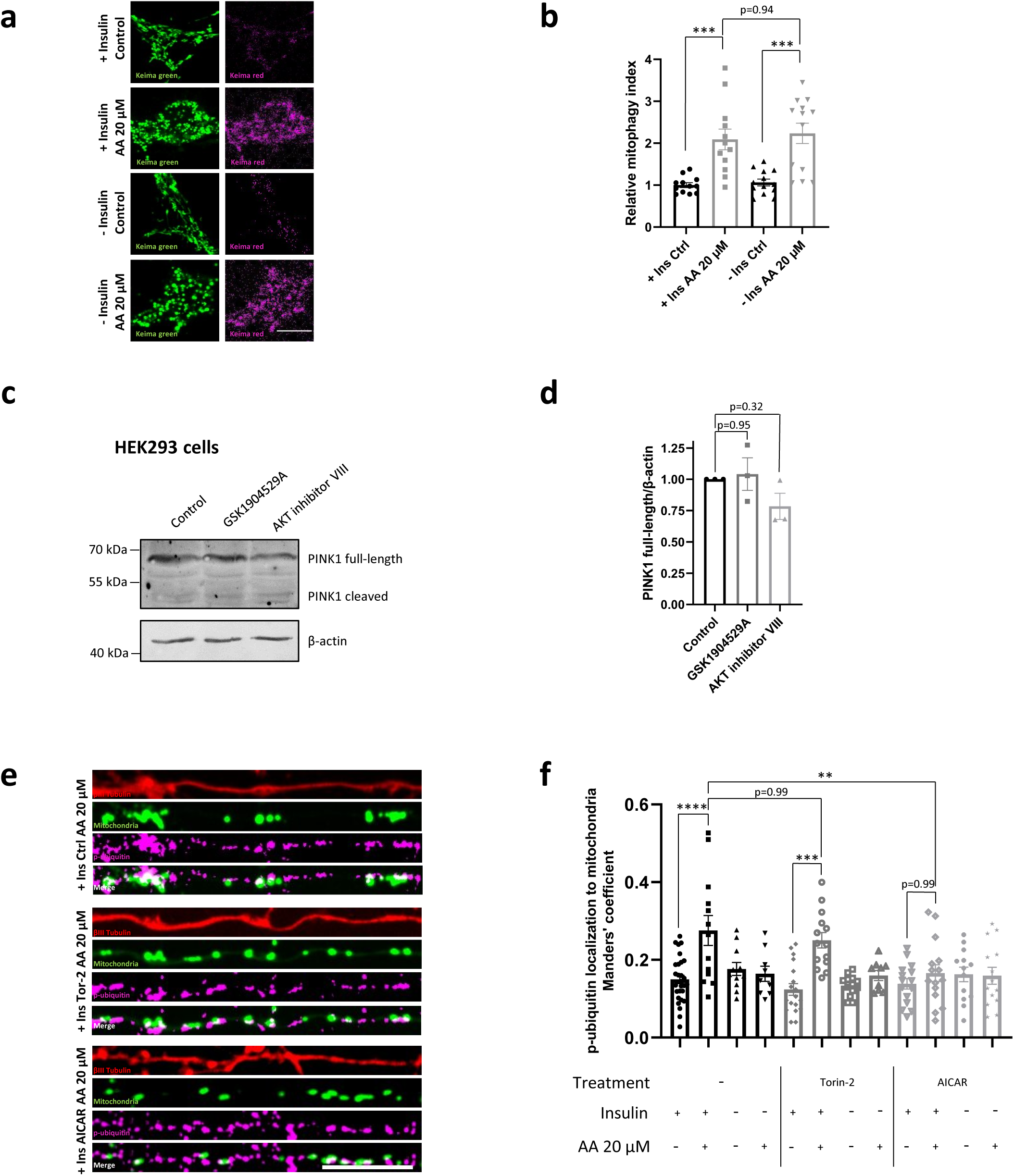
Insulin-regulated PINK1 expression is neuron-specific and mTOR-independent. **a** Representative images of neurons overexpressing the pH-sensitive fluorophore mito-mKeima cultured in the presence or absence of insulin overnight prior to treatment with or without 20 µM AA. **b** Quantification of the relative mitophagy index in neurons as in **a**. One-way ANOVA followed by Tukey’s post hoc test; n = 12-13; p < 0.001 (***). **c** Representative immunoblot image of HEK293 cells cultured with or without the IR inhibitor GSK1904529A (1 µM, 2 h) and the AKT inhibitor VIII (10 µM, 2 h), respectively. **d** Quantification of the full-length PINK1 protein bands normalized to the β-actin signal as in **c**. **e** Representative images of neurites overexpressing mito-meGFP cultured in the presence of insulin as well as Torin-2 (10 nM, 30 min) or AICAR (1 mM, 2 h) prior to treatment with 20 µM AA and stained with an antibody against phospho-ubiquitin (S65) and βIII tubulin. **f** Quantification of phospho-ubiquitin (S65) localization to mitochondria using the Manders’ colocalization coefficient. One-way ANOVA followed by Tukey’s post hoc test; n = 12-19; p < 0.01 (**), p < 0.001 (***), p < 0.0001 (****). All data are expressed as mean±SEM. Data points represent biological replicates (**d**) or single cells coming from ≥3 biological replicates (**b** and **f**). Scale bars, 10 µm.

**Extended Data Fig. 7 corresponding to Fig. 7.**
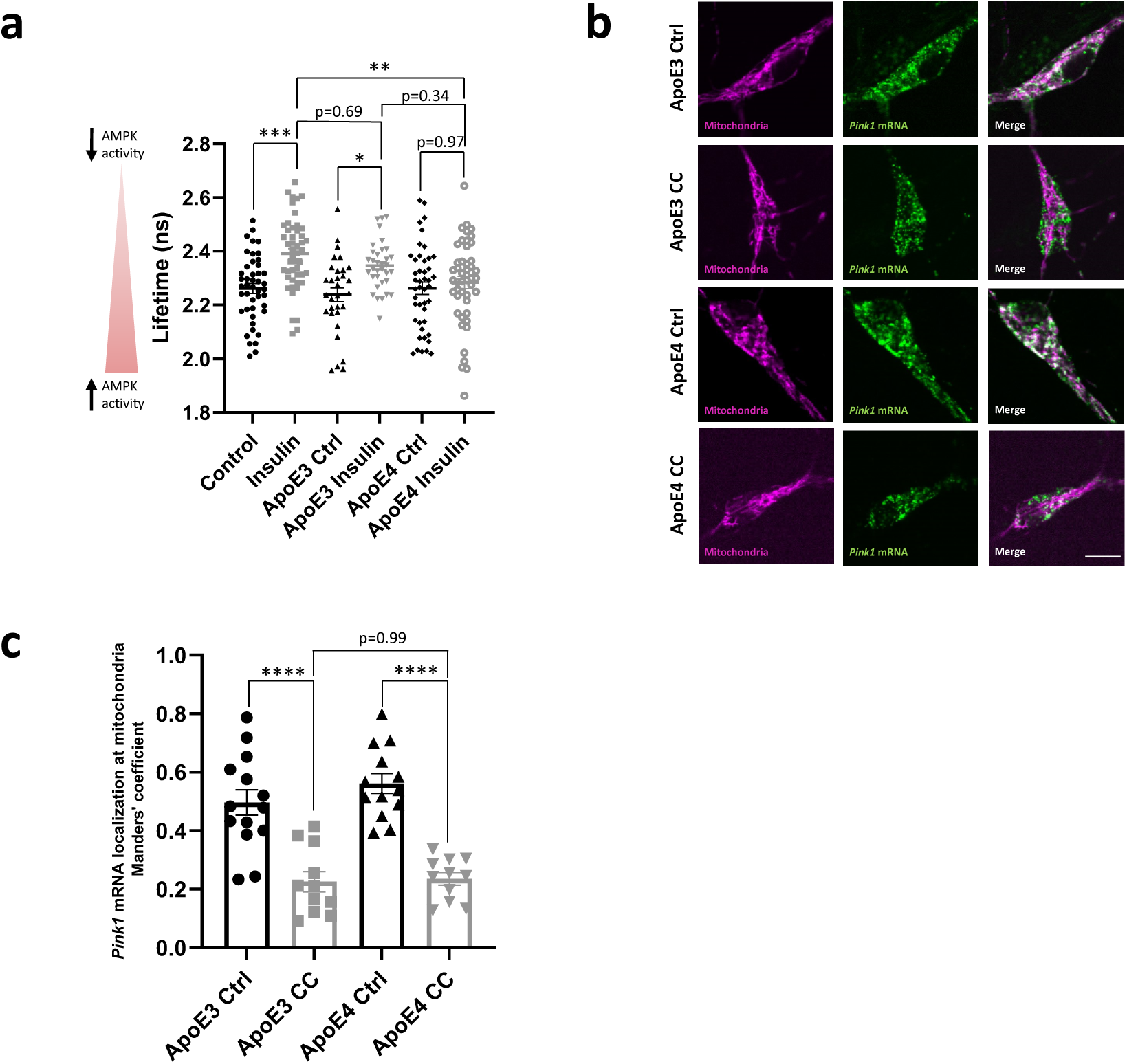
ApoE4 inhibits insulin-but not CC-mediated effects. **a** Quantification of fluorescent lifetime imaging of the FRET-based AMPK activity sensor in neurons treated with and without insulin (500 nM, 1 h) in the presence of ApoE3 (50 nM) and ApoE4 (50 nM) overnight, respectively. One-way ANOVA followed by Tukey’s post hoc test; n = 31-47; p < 0.05 (*), p < 0.01 (**), p < 0.001 (***). **b** Representative images of *Pink1* mRNA visualized by the MS2/PP7 split Venus method and mitoRaspberry with and without CC (20 µM, 2 h) treatment in the presence of ApoE3 (50 nM) and ApoE4 (50 nM) overnight, respectively. **c** Quantification of the Manders’ colocalization coefficient for the overlap between the *Pink1* mRNA and mitochondrial channel as in **b**. One-way ANOVA followed by Tukey’s post hoc test; n = 11-14; p < 0.0001 (****). All data are expressed as mean±SEM. All data points represent single cells coming from ≥2 biological replicates. Scale bars, 10 µm.

